# A minimal mathematical model for polarity establishment and centralspindlin-independent cytokinesis

**DOI:** 10.1101/2024.08.07.607072

**Authors:** Ondrej Maxian, Katrina M Longhini, Michael Glotzer

**Affiliations:** Department of Molecular Genetics and Cell Biology, University of Chicago, Chicago, IL 60637; Institute for Biophysical Dynamics, University of Chicago, Chicago, IL 60637

## Abstract

Cell polarization and cytokinesis are fundamental processes in organismal development. In the *Caenorhabditis elegans* model system, both processes are partially driven by a local inhibition of contractility at the cell poles. Polarization is triggered by centrosome-directed, local inhibition of cortical contractility at what becomes the posterior pole, while cytokinesis occurs when spindle elongation-dependent astral relaxation combines with centralspindlin-induced accumulation of myosin at the cell equator. During both processes, Aurora A (AIR-1) kinase is activated on centrosomes and diffuses to the cortex where it inhibits the guanine nucleotide exchange factor (GEF) ECT-2, attenuating RHO-1 activation and actomyosin-based contractility. While these biochemical processes have been characterized experimentally, a quantitative understanding of how this circuit drives cortical dynamics in polarization and cytokinesis is still lacking. Here, we construct a mathematical model to test whether a minimal set of well characterized, essential elements are necessary and sufficient to explain the spatiotemporal dynamics of AIR-1, ECT-2, and myosin during polarization and cytokinesis of *C. elegans*. We show that robust establishment of polarity can be obtained in response to a weak AIR-1 signal, and demonstrate the relevance of rapid ECT-2 exchange and a persistent AIR-1 cue during polarization. In the minimal model, the rapid turnover of ECT-2 causes a quasi-steady response to changes in the AIR-1 signal, and transient asymmetries are not self-sustaining. After tuning the model for polarization, we demonstrate that it can accurately predict ECT-2 accumulation during cytokinesis, suggesting a quantitative similarity between the two processes.

## 1. Introduction

Polarization and cytokinesis are fundamental processes in organismal development and physiology (Dewey et al., 2015). Cell polarization is encoded by asymmetric distributions of protein molecules, which are shaped by local regulation of binding and diffusion, and especially active transport by cortical flows (Munro et al., 2004; Mogilner et al., 2012; Lang and Munro, 2017). Likewise, cytokinesis involves the formation and constriction of the actomyosin ring in the cell mid-plane, a process which is driven by a balance of contractility, flow, and membrane mechanics (White and Borisy, 1983; White, 1985; Glotzer, 2005). The two processes can be biochemically and mechanically connected, as cell polarity can regulate spindle positioning, which controls the site of contractile ring assembly and, consequently, the division plane (Grill et al., 2001; Maddox et al., 2007, 2012; Davies et al., 2014).

The *C. elegans* zygote provides a powerful model in which to study both polarization and cytokinesis (Fig. 1). In *C. elegans*, as in many other animal cells, contractility in polarity establishment and cytokinesis is mediated by the GTPase RHO-1, which activates myosin through its effector Rho kinase. RHO-1 transitions between an active (GTP) state and inactive (GDP) state via interactions with the RhoGEF ECT-2 and RhoGAP RGA-3/4, which activate and inactivate RHO-1, respectively (Fig. 2(a); Michaux et al. (2018); Basant and Glotzer (2018)). Contractility in polarization and cytokinesis relies on modulation of this circuit. During polarization, contractility is activated by a nematode specific protein, NOP-1 which appears to globally activate ECT-2 (Tse et al., 2012). Contractility is inhibited by centrosomes at the position of sperm entry, establishing the posterior pole and triggering anterior-directed cortical flows that facilitate the segregation of anterior and posterior PAR proteins into distinct domains (Goldstein and Hird, 1996; Munro et al., 2004; Lang and Munro, 2017; Gross et al., 2019). During cytokinesis, the centralspindlin complex accumulates at the mid-plane of the spindle, where it activates ECT-2 and RHO-1, thus promoting contractility (Glotzer, 2005; Basant and Glotzer, 2018). This pathway combines with a second pathway similar to polarization, i.e. activation of RHO-1 spatially modulated by the centrosomes (Tse et al., 2012). Given the positions of the separated centrosomes, this also biases contractility to the cell equator (White, 1985; Werner et al., 2007; Loria et al., 2012). The goal of this study is to use mathematical models to determine whether the known regulatory pathways are sufficient to explain these highly stereotyped behaviors.

**Figure 1:**
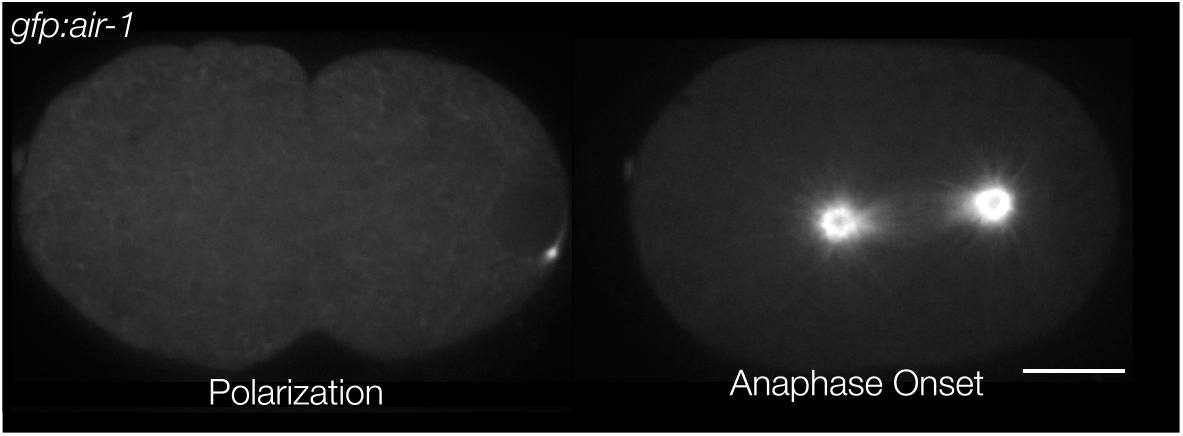
Aurora A (AIR-1) accumulation during polarization and cytokinesis in *C. elegans* embryos. AIR-1, which locally inhibits contractility, is enriched at the centrosomes. Anterior is positioned to the left, and the scale bar is 10 *µ*m.

**Figure 2:**
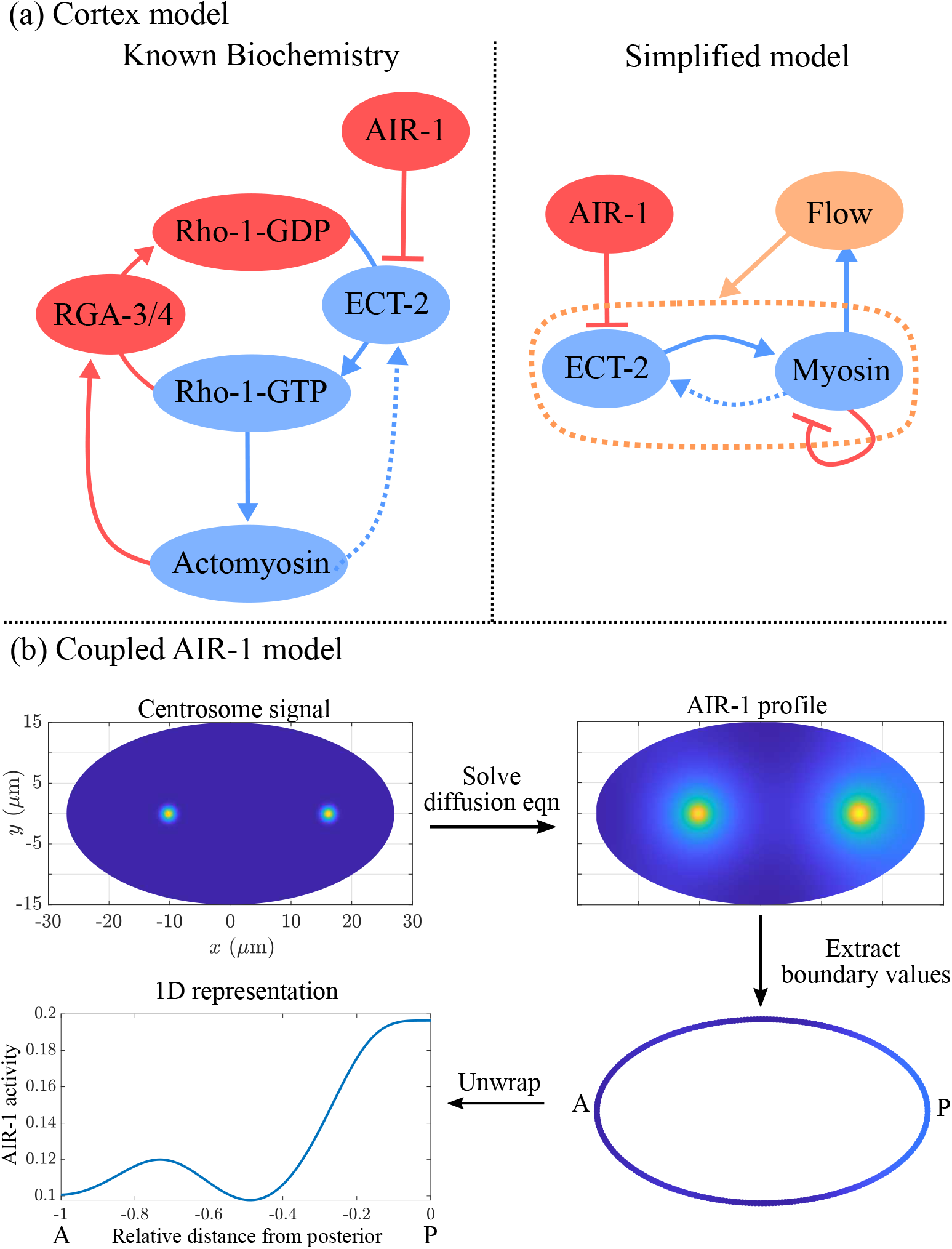
Modeling schematic for this study. (a) The model of the cortex, with the known biochemistry on the left and our simplified model on the right. This model takes the AIR-1 signal as input, and the equations are given in (1). (b) Procedure for determining the AIR-1 signal. Given centrosome positions (shown in cytokinesis in wild-type embryos), we solve for the AIR-1 profile on the cross section using equation (2), then formulate a one-dimensional AIR-1 profile by extracting the values on the boundary.

Recent studies have characterized the mechanism by which polarizing and dividing cells pattern the RHO-1 activation pathway. During polarization, Aurora A kinase (AIR-1) associates with the sperm centrosome, which is the sperm-derived structure that promotes polarity establishment (Hannak et al., 2001; Cowan and Hyman, 2004; Klinkert et al., 2019; Kapoor and Kotak, 2019; Zhao et al., 2019; Longhini and Glotzer, 2022). Recent work (Longhini and Glotzer, 2022) showed that AIR-1 impacts cortical dynamics by inhibiting ECT-2. Specifically, ECT-2 dissociates from the posterior membrane in an AIR-1-dependent manner, and it contains consensus sites for AIR-1 that are required for AIR-1 responsiveness. During polarization, ECT-2 exhibits posterior depletion and anterior enrichment, a pattern of accumulation that requires cortical myosin flows. A similar set of events occur upon anaphase onset, coincident with cytokinesis (Longhini and Glotzer, 2022). While the centrosomes have duplicated, matured, moved farther from the cortex (Fig. 1), and accumulated much more AIR-1 in cytokinesis, there remains a strong, ultra-sensitive dependence between the distance of the centrosome from the nearest cortical domain and the amount of cortical ECT-2 at that site; proximal centrosomes correlate with a reduction in cortical ECT-2 (Longhini and Glotzer, 2022).

While the qualitative mechanisms by which AIR-1, ECT-2, and myosin interact to generate contractility and flow have thus been well-characterized, how the hypothesized pathways could generate the quantitative patterns of myosin and ECT-2 accumulation during polarization and cytokinesis is still unclear. For instance, unlike the anterior PAR proteins, which have residence times on the order of one hundred seconds (Robin et al., 2014), ECT-2 cannot be strongly advected, as it exchanges rapidly between the cytoplasm and the cortex on timescales of a few seconds, appearing to preferentially accumulate on the cortex at myosin-enriched sites (Longhini and Glotzer, 2022). Consequently, it remains unknown whether a short residence time, preferential recruitment by myosin, and weak advection by cortical flows, could combine to generate the observed asymmetric accumulation of ECT-2 during polarization. More generally, it is not known if additional mechanisms are required to explain the pattern of ECT-2 accumulation during cytokinesis, as the centrosomes are so much further from the cortex at that stage.

In this study, we test whether a minimal set of interactions can explain the dynamics of ECT-2 and myosin during both polarity establishment and centralspindlin-independent cytokinesis. To do this, we construct a mathematical model that uses an AIR-1 signal, which diffuses from the centrosomes to the cortex, as an input to a continuum model of contractility (Fig. 2(b)), which is similar to those previously described (Michaux et al., 2018; Gross et al., 2019). We show that our model can explain the initial dynamics of polarization, similar to those observed in the absence of PAR proteins. Furthermore, the same model reproduces the patterns of ECT-2 accumulation observed during cytokinesis, thus demonstrating a quantitative similarity between the two processes.

## 2 Methods

In *C. elegans* embryos, the AIR-1 signal originates from the centrosomes, which are positioned in the interior of the three-dimensional cell (Fig. 1), while the contractile dynamics occur on the two-dimensional cell cortex (boundary). For our model, we consider a cross-section of the embryo, so that the AIR-1 dynamics occur in two dimensions, and the contractile dynamics occur on the one-dimensional boundary (Fig. 2(b)). The workflow is to first set the centrosome positions according to experimental data (Fig. 1), then solve a diffusion equation to obtain the AIR-1 profile at the cortex. This becomes an input to a set of one-dimensional reaction-diffusion-advection equations that treat the ECT-2/myosin relationship. In order to study how the cortex responds to the expected AIR-1 signals in polarization and cytokinesis, we assume that the centrosomes (and AIR-1 signal) are fixed (see Fig. S8 for a simulation that relaxes this assumption).

In this section, we first discuss the basic model of contractility, which follows previous work by others (Goehring et al., 2011; Gross et al., 2019). Building on this prior work, we seek to evaluate how a centrosomal AIR-1 signal stimulates cortical dynamics, and analyze the steady state ECT-2 accumulation during polarization and cytokinesis.

### 2.1 Basic model of contractility

At the cortex of the *C. elegans* zygote, AIR-1 inhibits accumulation of ECT-2 by increasing its dissociation rate through phosphorylation. The cortical pool of ECT-2 gains the ability to activate RHO-1, which activates myosin (Fig. 2(a)). Myosin feeds back on ECT-2 through advection by cortical flows, and there is nonlinear negative feedback of myosin accumulation (through RGA-3/4-dependent inactivation of RHO-1) (Michaux et al., 2018). To translate these dynamics into a simple model (Fig. 2(b)), we neglect the intermediary of RHO-1 and formulate a model with two variables, *E* (for ECT-2) and *M* (for myosin). In dimensional units, the equations we use are

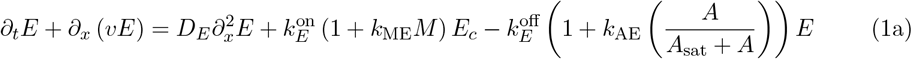

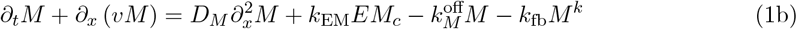

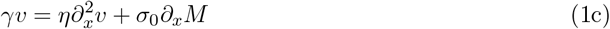

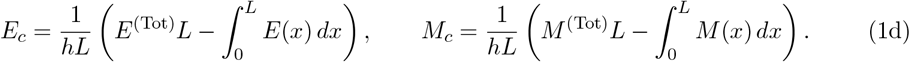

Similar to previous studies using these types of models (Goehring et al., 2011; Dawes and Munro, 2011; Kravtsova and Dawes, 2014; Gross et al., 2019), the model geometry is one-dimensional, and can be viewed as a one-dimensional slice of the cell cortex combined with a well-mixed cytoplasm (see Fig. 2(b)). Each species evolves by advection by cortical flows (terms *∂*_x_(*υE*) and *∂*_x_(*υM*) and), diffusion in the cortex (terms 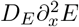, and 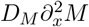), and binding/unbinding from the cortex. The binding rate is also proportional to the cytoplasmic concentration of each protein, defined in (1d), where *L* is the domain length, *h* is the cytoplasmic “thickness” (so that *hL* is the total area), and *E*^(Tot)^ is the concentration of ECT-2 when all of it is bound to the cortex (likewise for *M*) (Lang and Munro, 2022). Finally, the velocity equation (1c) expresses the balance of active stress (which we assume is proportional to myosin concentration) with viscous stress and frictional resistance (Mayer et al., 2010). For simplicity, along with many other components, we do not model the actin network, a topic for future study.

The binding/unbinding terms in the ECT-2 and myosin equations rely on the following major assumptions:

1. The variable *E* represents unphosphorylated, active ECT-2 bound to the cortex. We do not consider phosphorylation of ECT-2 in the cytoplasm, and instead assume that the effective ECT-2 binding rate (which we fit to experimental data) represents the binding rate of *unphosphorylated* ECT-2. Consistent with this assumption, the negative flux in ECT-2 represents the combined rate of unbinding and phosphorylation, with the latter being proportional to the AIR-1 concentration, except at high AIR-1 when it saturates (see Appendix A). Likewise, we assume that activation of ECT-2 by NOP-1 occurs uniformly throughout the cortex (Tse et al., 2012).
2. It was previously shown that ECT-2 has a small residence time at the cortex (on the order of a few seconds) (Longhini and Glotzer, 2022). Under these conditions, we show in Fig. S4 that the direct transport of ECT-2 by flows contributes negligibly to its steady state profile. It was previously speculated that ECT-2 could be effectively “transported” by associating with other components that are more stably bound to the cortex (Longhini and Glotzer, 2022). We incorporate this assumption into our model by assuming recruitment of ECT-2 by a species which is advected by cortical flows. For simplicity, in our equations we assume that the concentration of this species is equal to that of myosin, thus giving the *k*_ME_*ME*_*c*_ term in the ECT-2 equation (1a). In Fig. S7, we show that explicitly introducing a third species into the equations which recruits ECT-2 gives similar patterns during polarization.
3. Based on previous work which demonstrated an important role for RhoGAP in setting the size of the anterior domain in polarizing embryos (Schonegg et al., 2007), combined with other work showing a nonlinear relationship between RhoGAP activity and Rho/myosin accumulation (Nishikawa et al., 2017; Michaux et al., 2018), we postulate nonlinear negative feedback in the myosin kinetics, with an inactivation rate proportional to *M*^*k*^. As long as *k >* 1, this term provides a way of controlling potential instabilities that arise in the simple active gel model (1c) (Nishikawa et al., 2017). In the main text, we present results using *k* = 2, but in Fig. S5 we show that model predictions are similar when *k* = 3 instead.

#### 2.1.1 Parameter estimation

In Appendix B, we convert the model equations to dimensionless form, then fit the parameters using a combination of direct experimental measurements and inference based on other experimental data. The overall flow of the parameter fitting process goes as follows:

1. We first assign values to the diffusivities and unbinding rates of each component that come from direct experimental measurements (Gross et al., 2019; Goehring et al., 2011; Michaux et al., 2018).
2. Using previously-imaged embryos with myosin and ECT-2 markers (Longhini and Glotzer, 2022, Fig. 1), we measure the effective myosin and ECT-2 profiles during pseudo-cleavage (Fig. S3(a–c)), which we treat as a quasi-steady state.
3. Based on the quasi-steady myosin profile, we fit the velocity parameters in (1c) to match bulk flow speeds in wild-type embryos (which are typically at most 10 *µ*m/min) (Gross et al., 2019) (Fig. S3(d)).
4. To fit the parameters in the myosin equation (1b), we impose the measured ECT-2 profile and adjust *k*_fb_ and *k*_EM_ until we match the experimentally-measured myosin profile (Fig. S3(e)).
5. We use previous measurements in myosin depleted embryos (Longhini and Glotzer, 2022, Fig. 3A) to infer how AIR-1 impacts ECT-2 in the absence of myosin (i.e., to fit *k*_AE_) (see Fig. S4(a)).
6. With all other parameters fixed, we increase the rate *k*_ME_ at which myosin (or a species associated with myosin) recruits ECT-2 until we reach the boundary of contractile instabilities. We choose a value for *k*_ME_ that sits near the boundary between the stable and unstable regime, without giving unstable behavior (Fig. S4). The result of this parameter fitting is that about 40% of the ECT-2 that binds to the cortex is recruited by myosin-associated species.

**Figure 3:**
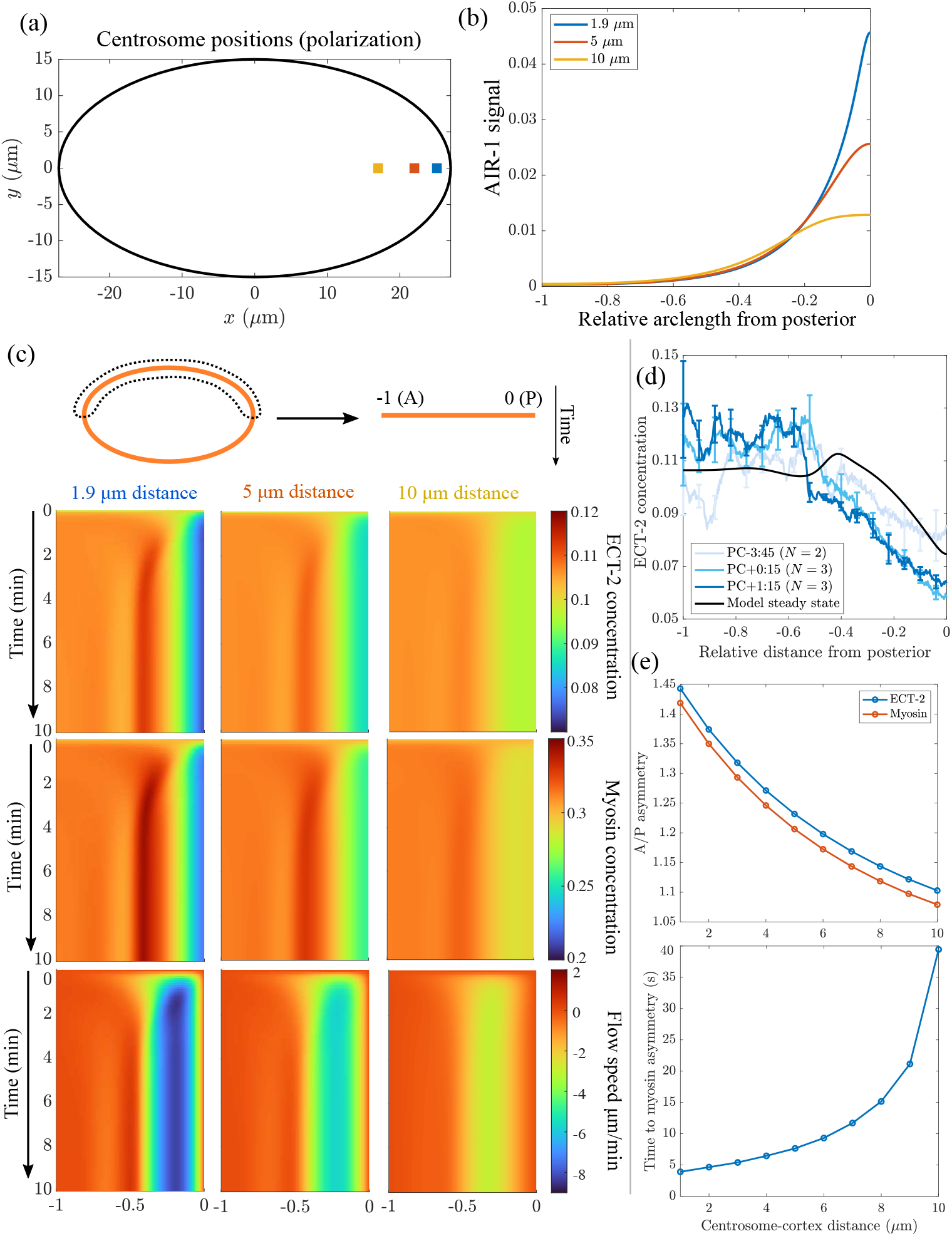
Centrosome locations set polarization dynamics. (a) The location of the centrosomes in our polarization simulations; we position both centrosomes 1.9, 5, and 10 *µ*m away from the cell boundary. (b) The resulting AIR-1 signals along the cell perimeter. (c) The dynamics of polarization, starting from the uniform state, with the computed AIR-1 signals. We show the ECT-2 concentration (top), myosin concentration (middle), and flow speed (bottom) in pseudo-kymographs, with time on the *y* axis, and the A/P axis on the *x* axis (A at left and P at right). (d) The steady state ECT-2 concentration in the model with centrosome-cortex distance of 1.9 *µ*m, compared to experimental ECT-2 profiles in polarizing embryos. See Appendix B.3.2 for data extraction procedure; the profiles are averaged over 2:30 (m:ss) time intervals, with times in the legend being the middle of the interval (PC = pseudocleavage furrow formation). (e) The A/P asymmetry in myosin and ECT-2 after 10 mins (top) and the time for symmetry breaking (bottom), both as a function of the centrosome-cortex distance.

While it is tempting to equate the model’s myosin-driven, contractile instabilities with pulsatile RHO-1/myosin excitability (Nishikawa et al., 2017), the latter are actually myosin-independent (Michaux et al., 2018), which illustrates that the wild-type embryo does not sit in the model’s fully unstable regime. In fact, the overall speed of bulk flows does not depend on pulsatility of RHO-1 (Michaux et al., 2018, Fig. 7), affording a biological justification for modeling the simpler case in which these well-characterized pulses of RHO-1 activation are absent (or, more precisely, the case where these randomly-positioned pulses are averaged over many cross sections to yield a steady signal). It was recently shown that knockdown of RHO-1’s downstream effectors could cause contractile instabilities, which implies that the wild-type *C. elegans* embryo likely sits near, but not within, the unstable regime (Yao et al., 2022). In this way, a comparison can be drawn to the *Xenopus* model system, where normally quiescent oocytes can exhibit excitable dynamics through over-expression of ECT-2 and RGA (Michaud et al., 2022; Chen et al., 2024).

As described further in Appendix B, we use a standard method of lines discretization to solve (1). At each time step, we first solve for the cortical flow speeds *v*, which we use in a first-order upwind finite volume scheme for the advection terms. The diffusion terms are treated implicitly using the backward Euler method.

### 2.2 Coupled model of AIR-1

As an input to the cortex model, we solve for the AIR-1 profile on the boundary (the cortex) of a two-dimensional embryo cross section by specifying the position of the centrosome(s) and solving a diffusion equation in the embryo interior. Letting *a*(***x***) be the concentration of AIR-1 in the two-dimensional embryo cross-section, its diffusion in the cytoplasm is described by

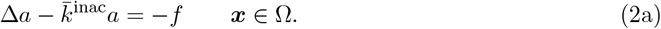

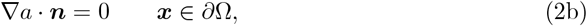

where (2a) is the diffusion equation for the concentration and (2b) is a no-flux boundary condition through the boundary (here Ω represents the embryo area and *∂*Ω represents the boundary). The signal *f* (***x***) comes from the two centrosomes, which we model by Gaussian densities

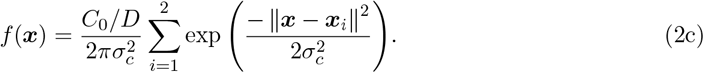

Here ***x***_*i*_ = (*x*_*i*_, 0) is the location of the *i*th centrosome, which changes depending on the experimental conditions. In addition to the centrosome location, the signal has two other parameters: *C*_0_*/D* is the strength of the cue (the integral of *f* (***x***) over the entire embryo cross-section, normalized by the cytoplasmic diffusion coefficient *D*), and 2*σ*_*c*_ is roughly the centrosome “size.” For cytokinesis, the centrosomes have radius about 1.4 *µ*m, so we set *σ*_*c*_ = 0.7 *µ*m. In polarization, the centrosomes have radius about 0.35 *µ*m, so we set *σ*_*c*_ = 0.175 *µ*m (Decker et al., 2011, Fig. 1C). The absolute signal strength *C*_0_*/D* is arbitrary, but ratios between cytokinesis and polarization are well defined. Consequently, we set *C*_0_*/D* = 1 for cytokinesis and *C*_0_*/D* = 1*/*32 for polarization, according to our experimental measurements (Fig. S2).

The diffusion equation (2a) also contains a basal rate of inactivation of AIR-1 (phosphatase activity). This introduces another parameter which is the inactivation rate relative to the diffusion coefficient in the cytoplasm (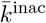, units *µ*m^*−*2^). As shown in Appendix A, low levels of phosphatase activity give high global AIR-1 levels, which translate to low ECT-2 levels everywhere. Such levels were shown to block pseudocleavage in centralspindlin-independent cytokinesis, due to low contractility (Afshar et al., 2010; Kotak et al., 2016). We choose the phosphatase activity level such that centrosomes close to the posterior pole (in polarization) negligibly impact the anterior domain (*<* 1% of the posterior concentration); see Fig. S1.

Once the parameters are set, we use a standard first-order finite element method to solve (2a). In brief, the elliptical domain of the embryo is meshed into nodes and triangles (Persson and Strang, 2004), and the finite element matrix equation becomes 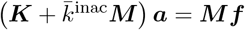, where ***M*** is the mass matrix and ***K*** is the stiffness matrix for finite elements, which are assembled using standard techniques (Gockenbach, 2006, c. 7). Solving for *a*(***x***) everywhere gives a profile on the embryo perimeter (cortex), which we substitute into (1) as the one-dimensional profile *A*(*x*), using linear interpolation to map from the (irregular) finite element boundary nodes to a regular grid. Once it reaches the cortex, AIR-1 inhibits ECT-2 by increasing its cortical dissociation rate, in correspondence with experimental data in myosin-depleted embryos (Longhini and Glotzer, 2022, Fig. 3A).

## 3 Results

We now use the model to explore the quantitative similarities between polarization and cytokinesis. As discussed in Section 2 and Appendix B, some of the parameters are constrained using measurements from the polarized wild-type embryo. As such, the statements we make about wild-type polarization are shaped by existing data. Changing parameters, such as the centrosome-cortex distance and ECT-2 residence time, will demonstrate the importance of the centrosome position and relevance of rapid ECT-2 exchange during polarization. Perhaps more important is the model’s direct extrapolation to cytokinesis, which concludes this section. Without changing any parameters, we demonstrate that the model can also explain the observed ECT-2 accumulation patterns during cytokinesis (Longhini and Glotzer, 2022), which are cued by larger centrosomes with 30-fold more AIR-1 than in polarization (Fig. 1).

### 3.1 The centrosome distance determines the strength of polarization

During cell polarization, the newly-duplicated centrosomes typically sit very close to the posterior cortex (about 1.9 *µ*m away) and are small (approximate radius of 0.4 *µ*m), as they have not yet accumulated large amounts of pericentrosomal material (Bienkowska and Cowan, 2012; Decker et al., 2011). To explore the effect of centrosome distance and location, we position centrosomes at a distance 1.9, 5, and 10 *µ*m from the posterior pole, and measure the resulting AIR-1 signal by solving the diffusion equation (2) on the embryo cross section and extracting boundary values (see schematics in Figs. 2(b) and 3(a)). As expected, there is a significant decrease in the AIR-1 signal as the distance between the centrosomes and the cell boundary increases. As compared to a 1.9 *µ*m distance, centrosomes 5 *µ*m from the cortex exhibit a decrease of 50% in AIR-1, and those 10 *µ*m from the cortex exhibit a further decrease of 50% (Fig. 3(b)).

Using these profiles of AIR-1 activity, we run the cortex model forward in time to reach a steady state for polarization. For reference, in the absence of PAR proteins, the centrosomal signal induces a transient clearing of myosin from the posterior pole, and the myosin profile reverts back to a uniform state after the centrosomes move towards the cell equator (Gross et al., 2019, Fig. 2E). Figure 3(c) shows that our simulations reproduce the initial smaller-scale clearing of both myosin and ECT-2. Notably, the steady state ECT-2 accumulation in our simulation (which is steady because the centrosome positions are fixed) matches experimental data from early establishment phase (four minutes before pseudo-cleavage), but diverges from the quasi-steady state that emerges once polarity is established (Fig. 3(d)), perhaps reflecting the influence of PAR proteins in the network (Gross et al., 2019). The predicted domain of ECT-2 clearance comprises about 30% of the half-perimeter (∼20 *µ*m on either side of the pole), which is in good agreement with experimental observations in PAR mutants (Gross et al., 2019, Fig. 2E), suggesting that the model can effectively capture the transient posterior clearing induced by the AIR-1 signal when PAR proteins are absent. In the model, changing the distance between the centrosomes and the cortex affects the quantitative values of the asymmetries and the time to reach them (Fig. 3(e)), but not the myosin peak location. As shown in the kymographs in Fig. 3(c), the peak location, which correlates with the position of the pseudocleavage furrow (Reymann et al., 2016), is roughly the same across all conditions, since it is controlled by the hydrodynamic length-scale (20% of the half-perimeter). Similar to previous observations (Bienkowska and Cowan, 2012, Fig. 3F), the time for “symmetry breaking,” defined as a local (5%) clearance in myosin from the posterior pole, is ten-fold higher when centrosomes are 10 *µ*m away from the cortex than when they are 1 *µ*m away. However, our model predicts an exponential scaling of the time to symmetry breaking at very large distances; this does not match the linear trend up to 10 *µ*m that was previously reported (Bienkowska and Cowan, 2012, Fig. 3F).

### 3.2 Rapid exchange and indirect recruitment

We have shown that long-range redistribution of ECT-2 is possible despite its short residence time. The driver of this redistribution is our assumption of indirect recruitment of ECT-2 by a longer-lived species which is advected by cortical flows. To explore how this assumption influences polarization kinetics, in Fig. 4 (left two columns) we consider two alternative models for how ECT-2 might segregate during polarization. In the first model, we remove recruitment of ECT-2 (by setting *k*_ME_ = 0 in (1a)), and observe at most 10% clearing at the posterior. Indeed, with realistic flow speeds of at least 5 *µ*m/min (Gross et al., 2019), molecules with residence time of 5 s can move at most 0.4 *µ*m, which is less than 0.5% of the embryo perimeter, indicating that cortical flows alone are insufficient to reproduce typical ECT-2 clearance levels.

**Figure 4:**
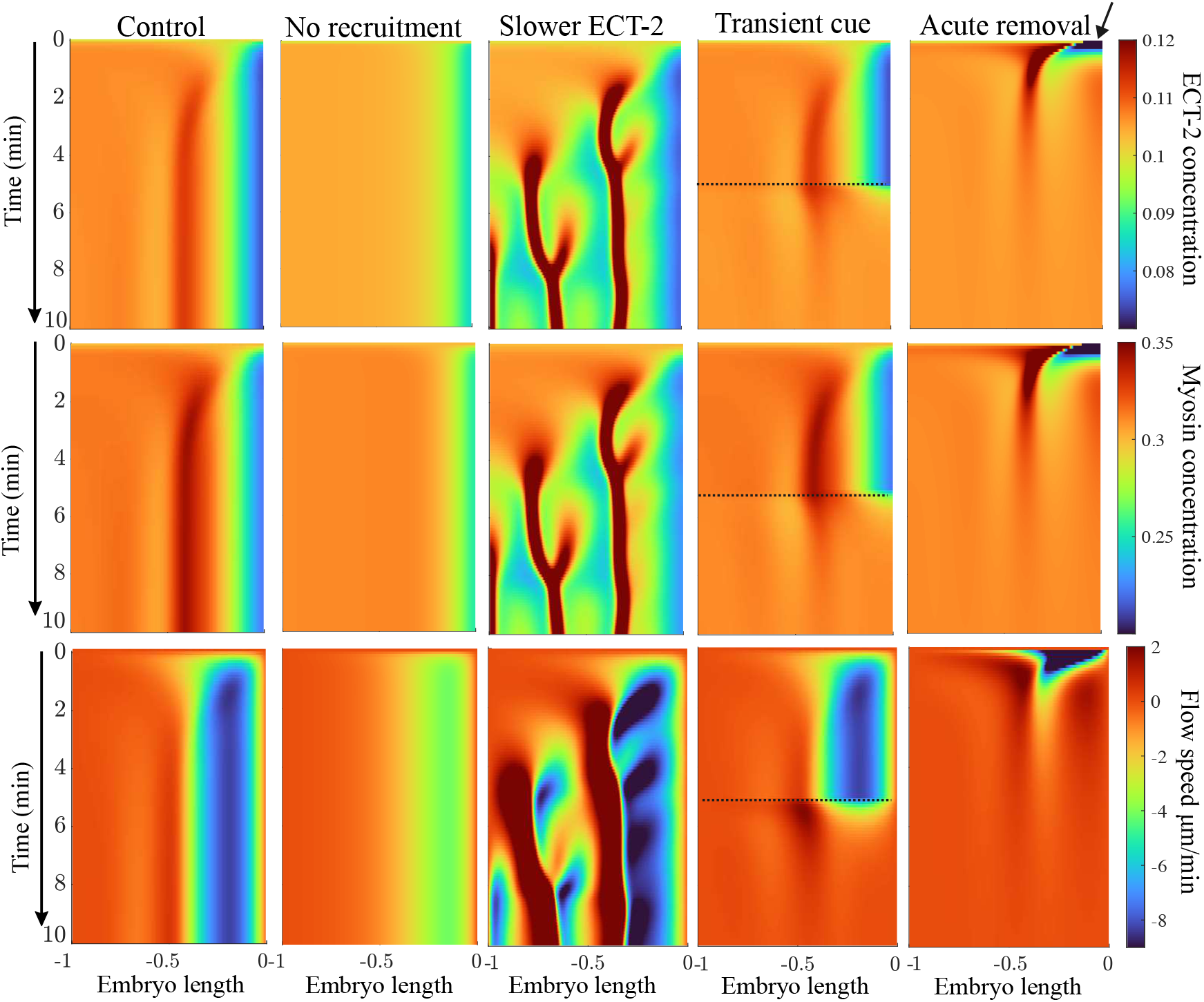
Adjusting conditions in polarization. Left panel: control parameters. Second from left: simulating the same parameter set without myosin-mediated recruitment of ECT-2 (*k*_ME_ = 0). Middle panel: adjusting parameters so that, rather than be recruited by myosin, ECT-2 has a four-fold longer residence time and is only advected. Second from right: simulating the case where the AIR-1 cue is present for five minutes (up to the dotted lines), after which we remove the AIR-1 signal and follow relaxation. Right panel: Simulating the case of no AIR-1 signal, but an initial condition in which ECT-2 and myosin concentrations are set to zero in a posterior region equal to 10% of the embryo (arrow).

An alternative model, albeit inconsistent with measured rates of ECT-2 exchange, is one where ECT-2 stably associates with longer-lived components on the cortex (for example through oligomerization (Illukkumbura et al., 2023)), so that the lifetime of ECT-2 is four times longer on the cortex. In this case, we observe contractile instabilities not seen in wild type embryos. As in control conditions, a local maximum in ECT-2 initially forms near ∼30% embryo length. Because of the increased residence time, however, ECT-2 is also advected *from the anterior* into the peak, which creates a local minimum in myosin and drives the formation of a second, posterior peak. Thus the rapid exchange of ECT-2 is important for stabilizing unidirectional flows during polarization. This outcome represents an experimentally-testable prediction, whereby a version of ECT-2 with more stable binding kinetics should produce contractile instabilities.

### 3.3 Assessing alternative polarity cues

The contractility cue that drives polarization establishment has been suggested to result from dynein-dependent removal of myosin (Chapa-Y-Lazo et al., 2020). Although experiments have shown that polarity establishment is dynein-independent (Longhini and Glotzer, 2022), we nevertheless considered whether a transient reduction in contractility would be sufficient to trigger polarity establishment. In particular, we perform two simulations: one with the AIR-1 cue active for five minutes, and a second that lacks an AIR-1 cue but in which we acutely remove myosin and ECT-2 at the posterior pole. The resulting dynamics over ten minutes are shown in Fig. 4. In both cases, transient cues or initial conditions ultimately relax to a steady state in which ECT-2 and myosin are nearly uniform and flow speeds are near zero.

The main difference between the transient AIR-1 cue and acute removal of myosin is in the intermediate dynamics. For a transient cue, the flow starts at zero, and steadily increases over the first minute, peaking with a flow speed of 8 *µ*m/min. Turning off the cue causes the flow speeds to slow below 2 *µ*m/min in less than one minute. By contrast, unloading myosin and ECT-2 from the posterior at *t* = 0 triggers a similar set of flows towards the anterior, but the flows are maximal at *t* = 1 minute, and then steadily decrease over time. Experimental data in PAR mutants show flows that reach a maximum velocity almost immediately after polarity triggering, but the magnitude (5 *µ*m/min) of these flows persist throughout polarity establishment phase (3–4 mins) (Gross et al., 2019, Fig. 2G). Thus, this supports models in which AIR-1 triggers a local and persistent inhibition of contractility, similar to that reported previously (Gross et al., 2019).

The rapid recovery of ECT-2 and myosin in our transient cue simulation comes from the complete removal of the AIR-1 cue at *t* = 5 mins. Of course, the cue’s removal is more gradual *in vivo*, and corresponds to steady motion of the centrosomes away from the posterior cortex. In Fig. S8, we demonstrate that a much longer timescale of posterior recovery results when we model this (more realistic) case. The modeled recovery of myosin with relocalizing centrosomes compares well with previous results that measured myosin recovery in PAR-depleted embryos (Gross et al., 2019, Fig. 2E).

### 3.4 ECT-2 accumulation in cytokinesis

The asymmetric accumulation of ECT-2 on the cortex during cytokinesis is sensitive to the position of the two centrosomes (Longhini and Glotzer, 2022). A plot of ECT-2 accumulation as a function of distance from the anterior/posterior pole closest to the centrosome reveals an “S-shaped” curve; at short and long distances there is a plateau in the ECT-2 accumulation (Longhini and Glotzer, 2022, Fig. 7A). By contrast, for distances in the range 10–20 *µ*m, there is an ultra-sensitive dependence of the ECT-2 concentration on the proximity. Note that during cytokinesis, there are two mature centrosomes which contain significantly more AIR-1 (about 30 times, Fig. S2) than the immature centrosomes that trigger polarity establishment.

To test whether the model that accurately predicts polarization recapitulates the behavior of ECT-2 during cytokinesis, we model the behavior of ECT-2 and myosin using the centrosome positions measured previously (Longhini and Glotzer, 2022), and repeated in Fig. 5. We consider the case of wild-type embryos and three representative experimental conditions, two with asymmetric centrosome positions (*dhc-1* (RNAi) and *zyg-9* (b244)), and one (*par-2* (RNAi)) with symmetric centrosome positions. As in polarization, we solve the diffusion equation (2) on the embryo cross section with the given centrosome positions, then extract the AIR-1 signal (in arbitrary units) along the cell perimeter (Fig. S2). Despite a much (four-fold) stronger posterior AIR-1 signal in *dhc-1* (RNAi) embryos, the A/P asymmetry in ECT-2 accumulation only changes by about 25% compared to *zyg-1* (b244); in Appendix A, we use this observation to constrain the AIR-1 saturation level *A*_sat_ that appears in (1a).

**Figure 5:**
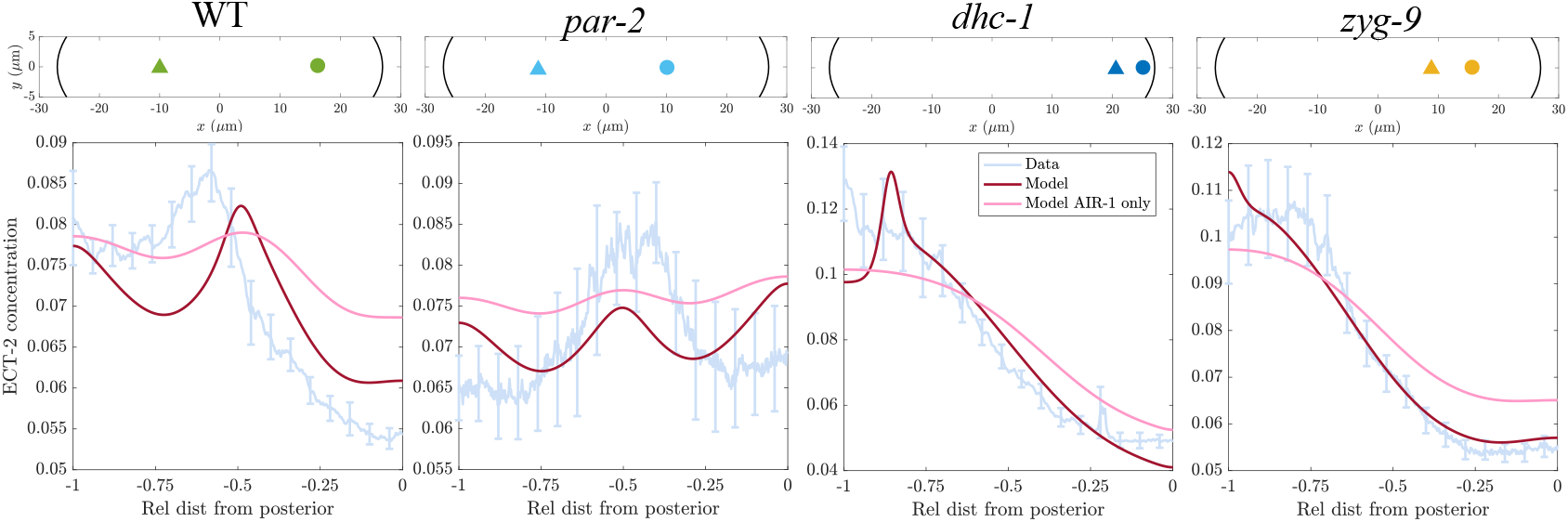
Extending the model to infer ECT-2 profiles in cytokinesis. We consider four experimental conditions from (Longhini and Glotzer, 2022): wild type (*N* = 10), *par-2* (RNAi) (*N* = 5), *dhc-1* (RNAi) (*N* = 10), and *zyg-9* (b244) (*N* = 9). For each condition, we show the average centrosome positions in the top panel. The corresponding bottom panel shows the experimental ECT-2 profile, averaged over a 50 s window beginning at cleavage furrow formation, compared with the steady state model result (dark red lines), and the model’s steady state when ECT-2 is only impacted by AIR-1 and not myosin (*k*_ME_ = 0 and *v* = 0 in (1a)).

We use the computed AIR-1 signals as inputs to the same cortex model (Fig. 2(a)) that we parameterized under polarization conditions. Without changing any of the parameters, we simulate each experimental condition to steady state, then compare the profile of ECT-2 accumulation with the experimental data taken over a 50 s window beginning at cleavage furrow formation (multiplied by a constant to match the mean model concentration). For the four centrosome positions simulated, the model quantitatively reproduces the ECT-2 accumulation pattern (Fig. 5, see Fig. S9 for AIR-1 depletion with wild-type centrosome positions). In wild-type and *par-2* (RNAi) embryos, the model qualitatively reproduces the observed profile, but tends to underestimate the A/P asymmetry. The shift in the central peak is likely due to centralspindlin-directed ECT-2 accumulation (Longhini and Glotzer, 2022), which is not accounted for in the model. In all experimental conditions, the fit to the data significantly degrades when we remove cortical flows and indirect recruitment (setting *v* = 0 and *k*_ME_ = 0 in (1a)). Thus, as during polarization, cortical flows amplify weaker AIR-1 signals. Indeed, as shown in Fig. S9, the predicted ECT-2 profile without myosin flows and indirect recruitment matches experimental data with partial myosin depletion, indicating that the model correctly accounts for both the response to AIR-1 signals and flow-based amplification (albeit imperfectly in some cases).

## 4 Discussion

Cell polarization in the *C. elegans* zygote is dependent on a centrosome-dependent signal that locally inhibits contractility. Similarly, cytokinesis can be influenced by aster positioning (White, 1985; Dechant and Glotzer, 2003; Severson and Bowerman, 2003; Munro et al., 2004). Only recently, however, has a better understanding of the nature of these cues emerged (Gross et al., 2019; Kapoor and Kotak, 2019; Klinkert et al., 2019; Zhao et al., 2019; Longhini and Glotzer, 2022; Manzi et al., 2024). In particular, a recent study (Longhini and Glotzer, 2022) showed that Aurora A kinase, AIR-1, which accumulates and is presumably activated at centrosomes, causes inhibition of the RhoGEF ECT-2 (and, consequently, RHO-1 and myosin) at the proximal cortex. The goal of this study was to determine whether a minimal mathematical model could explain experimental observations in both polarization and cytokinesis.

We introduced a hybrid two- and one-dimensional model (Fig. 2), where the centrosome positions are obtained from experimental data, and used to obtain a cross sectional profile of AIR-1 activation. Assuming diffusion of AIR-1 to the cortex (boundary of the cross section), we then obtained the profile of AIR-1 as an input to a model of cortical contractility. This cortex model, which we defined on the cross sectional-boundary and consequently made one-dimensional, included negative feedback of AIR-1 on ECT-2 accumulation, and positive feedback of myosin on ECT-2 accumulation, both through advection and recruitment by advected species. The coupling of the AIR-1 and cortex models allowed the dynamics of the active cortex to be dictated by the positions of the centrosome(s). We constrained the model using experimental observations (Longhini and Glotzer, 2022) and assuming modest (at most 2-fold) effects on cortical ECT-2 by Aurora A (negatively) and myosin (positively). The parameters arising from these constraints placed the model in a regime which can amplify small signals without yielding to contractile instabilities (Fig. S4).

This minimal model recapitulated many of the pertinent observations from polarization and cytokinesis. It produced transient polarization (in the absence of PAR proteins) with the observed flow speeds and protein ratios, and revealed the need for a *persistent* AIR-1 cue over time to maintain a polarized state in the absence of PAR reorganization. The picture that emerged from our experiments and modeling is a dynamic ECT-2 molecule that rapidly exchanges with the cortex, being preferentially recruited by longer-lived, flow-coupled molecules to sustain polarization. In this way, it reproduces other examples from cell biology where stable configurations are mediated by transient interactions (Ladurner et al., 2016). The rapid exchange of ECT-2 and, to a lesser extent, myosin, gives the model a quasi-steady nature; changes in the distribution of AIR-1 rapidly establish a new steady state. These dynamics are consistent with previous experimental results (Longhini and Glotzer, 2022, Fig. 2A), which showed an acute response of cortical ECT-2 to spindle rocking during anaphase on a timescale of 10 s, and may help to explain the rapid repositioning of the cleavage furrow in response to spindle displacements (Rappaport, 1985).

It is instructive to contrast the role of ECT-2 in generating cortical flows to that of the anterior PAR proteins, specifically PAR-3. While it has long been known that reduced PAR-3 levels correlate with reductions in cortical flows, these changes have only recently been linked to the *residence time* of PAR-3 molecules on the membrane (Chang and Dickinson, 2022; Illukkumbura et al., 2023). In wild-type embryos, PAR-3 monomers (which have residence time less than 1 s) (Lang et al., 2024) oligomerize to stably bind the membrane (residence times on the order 100 s), which allows them to both create and be advected by cortical flows (Munro et al., 2004). Consequently, embryos with oligomerization-defective PAR-3 fail to polarize because of a lack of coupling to (weaker) cortical flows. Our analysis indicates that ECT-2 lives in a different part of the stability diagram of chemical-mechanical coupling. In wild type embryos, the ECT-2 exchange kinetics (as measured by FRAP) are on the order of a few seconds, yet large-scale cortical flows are generated. Consequently, the model predicted that a longer ECT-2 residence time would produce hypercontractility, specifically a counterflow from the anterior end of the cell that prevents proper polarization. This result represents an important prediction which can be tested experimentally. The differences in stability behavior might be due to the coupling of ECT-2 and PAR-3 to myosin: whereas the role of PAR-3 in generating flows is likely indirect, ECT-2 directly generates flows by activating Rho and myosin. Thus, more tunable control of contractility could be achieved by faster turnover rates in the latter case.

The question that underpins our work is how a persistent flow could affect the distribution of transiently-bound proteins, independent of the underlying biochemical circuit in which they operate. For PAR-3, the typical measured diffusivities (0.01 *µ*m^2^/s) and residence times (200 s) are insufficient to explain the measured asymmetries when only advection and diffusion are assumed to contribute to patterning (Illukkumbura et al., 2023, Fig. 6H). This problem only worsens with ECT-2, which is apparently patterned by myosin-mediated flows, despite having a residence time on the order of a few seconds (Longhini and Glotzer, 2022). Here, we found that introducing recruitment of ECT-2 by a longer-lived species (or any species that is advected by and colocalizes with myosin) could result in similar patterning as cortical flows. Given that we previously showed that ECT-2 only segregates when its PH domain is intact (Longhini and Glotzer, 2022, Fig. 5A), we speculate that myosin could advect factors that cause PIP2 to concentrate anteriorly (Scholze et al., 2018; Hirani et al., 2019; Nakayama et al., 2009) which would then contribute to ECT-2 recruitment. This model could explain how ECT-2 segregates while turning over rapidly.

Because the model parameters were tuned to match observations during polarization, it was most striking that the model also predicted patterns of cortical ECT-2 accumulation in cytokinesis across multiple experimental conditions solely by modifying centrosome size and positions (Dechant and Glotzer, 2003; Verbrugghe and White, 2007; Longhini and Glotzer, 2022). Similar to recent work on contractility during cytokinesis (Werner et al., 2024), we found that the accumulation of ECT-2 in different conditions could be better reproduced by incorporating mechanochemical feedback to amplify the AIR-1/ECT-2 signal. Historically, inhibition of cortical contractility by asters was thought to be microtubule-mediated, given their proximity to the cortex (Dechant and Glotzer, 2003; Motegi et al., 2006). Yet experimental evidence (Klinkert et al., 2019; Kapoor and Kotak, 2019; Zhao et al., 2019; Longhini and Glotzer, 2022) and this mathematical model indicate that centrosome position and embryo geometry (i.e., the ability for AIR-1 to diffuse from the centrosomes to the cortex) serve as the primary determinants of polar relaxation. However, as cortical interactions with astral microtubules frequently control spindle and hence centrosome positioning (Grill et al., 2001; Schaefer et al., 2000; Tame et al., 2014), microtubules nevertheless play a role in this process.

## Data availability

See the github repository https://github.com/omaxian/CElegansModel/ for code.

## A AIR-1 diffusion model

### A.1 Constraining level of AIR-1 phosphatase activity

To constrain the level of phosphatase activity (parameter 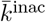 in (2)), we compute the profile of AIR-1 under polarization conditions (both centrosomes 1.9 *µ*m from the posterior pole), and plot the resulting AIR-1 signal in Fig. S1. When the phosphatase activity is low, the no flux boundary condition traps AIR-1 inside the embryo, and the relative difference between posterior and anterior is small. Increasing the phosphatase activity disproportionately lowers AIR-1 levels on the anterior pole (since it is much farther from the centrosomes). We assume that anterior levels of AIR-1 during polarization are at most 1% of posterior levels; this constrains 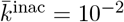.

**Figure S1:**
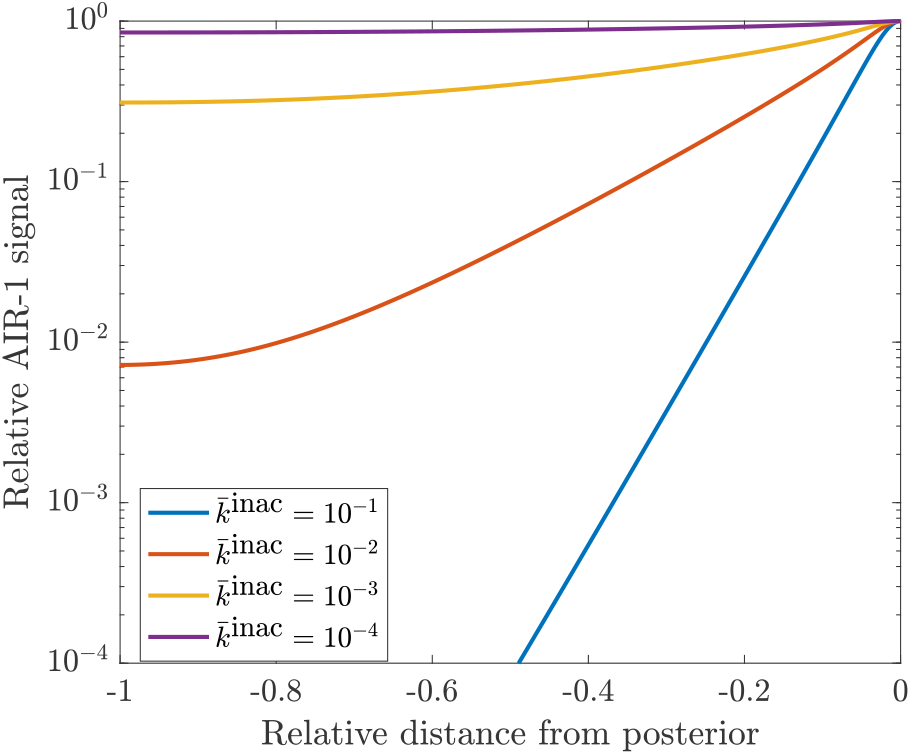
AIR-1 signal vs. distance from posterior under polarization conditions (blue squares in Fig. 3(a)). We vary the level of phosphatase activity 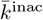 until the posterior level is less than 1% of the anterior level, settling on 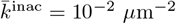.

To infer the profile of AIR-1 during cytokinesis for the four experimental conditions shown in Fig. 5, we use the corresponding centrosome positions (with four-fold larger centrosomes and 32-times more AIR-1 at each centrosome), set 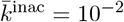 and solve (2). The resulting AIR-1 profile along the embryo perimeter is shown in Fig. S2. Despite large (four-fold) differences in posterior *zyg-9* and *dhc-1* AIR-1 signal, the A/P asymmetry between the two conditions increases by only 25% (Fig. 5). Because of this, we conclude that the saturation level of AIR-1 activity, *A*_sat_ in (1a), must lie near the *zyg-9* posterior levels. As such, we set *A*_sat_ = 0.25.

**Figure S2:**
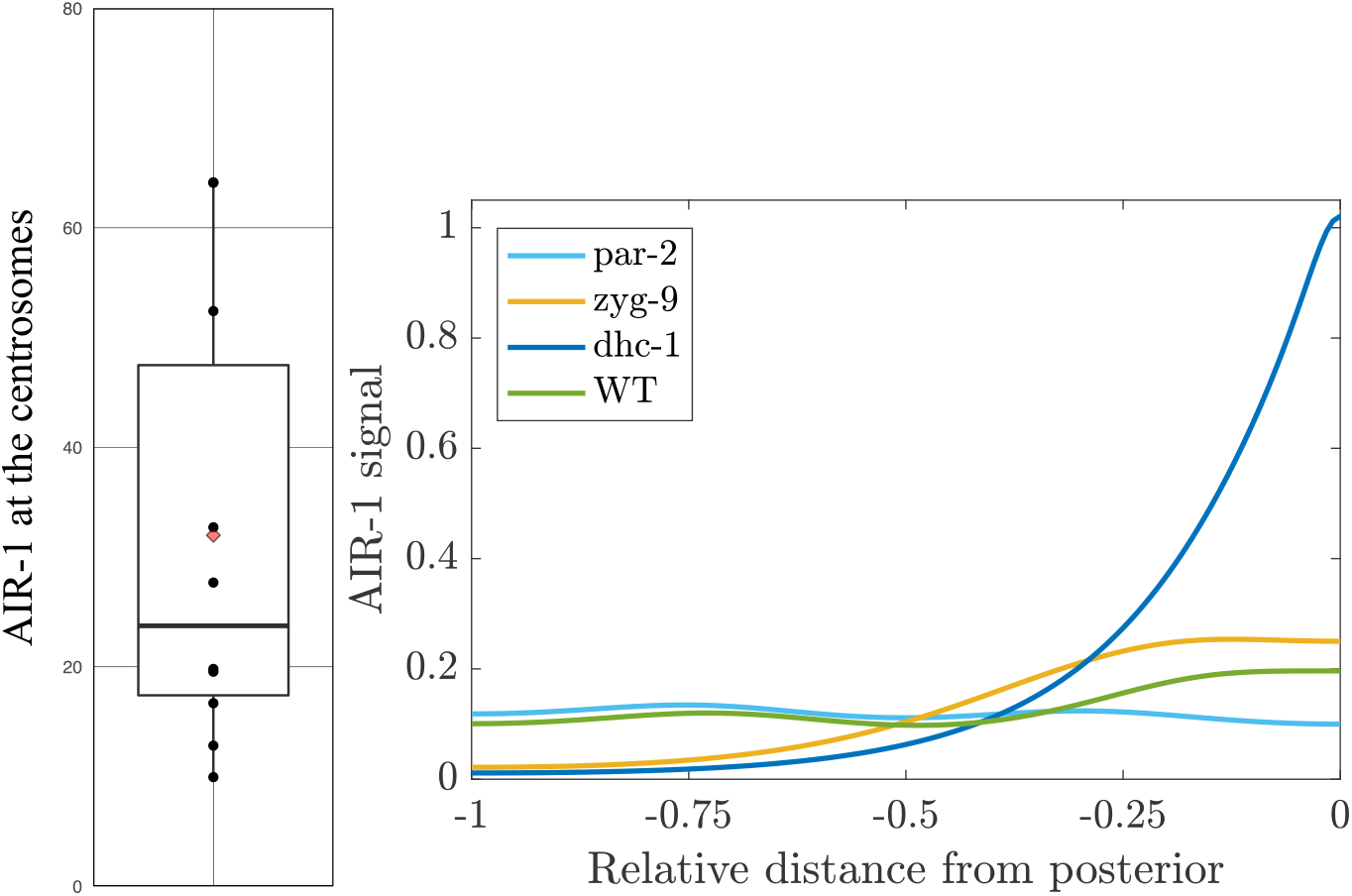
AIR-1 signal during cytokinesis. Left plot: the amount of AIR-1 at anaphase onset relative to polarization, which has mean 32 (*n* = 10). The black circles represent individual embryos and the red diamond is the mean. Right plot: The corresponding AIR-1 profile on the embryo boundary, which comes from the solution of the diffusion equation (2) on the embryo cross section (see Fig. 2(b)).

## B ECT-2/Myosin circuit

### B.1 Dimensionless form

Because absolute concentrations are unknown, it is easiest to assign values to unknown parameters when they are in dimensionless form. To do this, we non-dimensionalize (1) so that length is in units of the embryo perimeter *L*, time is in units of the bound myosin lifetime 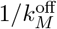, velocity is in units of 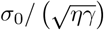 (Bois et al., 2011) *and concentration of species A* is in units of *A*^(Tot)^. This gives new dimensionless variables (denoted by hats)

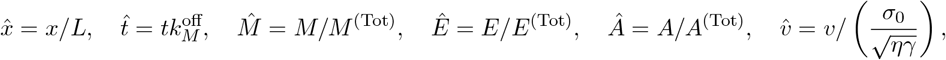

and a corresponding set of equations

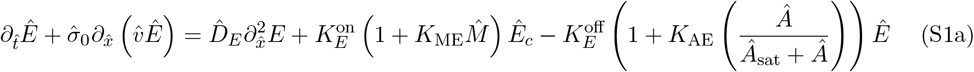

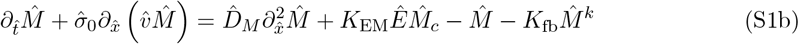

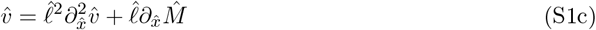

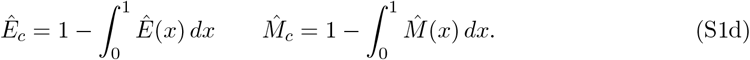

The conversion from dimensional to dimensionless form is given for all parameters in Table 1. Most of these conversions are straightforward, but there are some important parameters to highlight. In flow patterns, 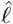 is a hydrodynamic length scale (scaled by domain perimeter) expressing the connectivity of the cortex; local disturbances in myosin will typically propagate at most a distance 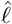. The parameter 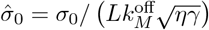 expresses the strength of the flows; the dimensional velocity in *µ*m/s can be extracted by taking 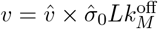.

**Table 1.** Relationship between dimensionless and dimensional parameters.

| Dimensionless parameter | Dimensional expression | Description |
| --- | --- | --- |
| $\hat{\sigma}_0$ | $\sigma_0 / (L k_M^{\text{off}} \sqrt{\eta \gamma})$ | Flow strength |
| $\hat{\ell}$ | $(\sqrt{\eta / \gamma}) / L$ | Hydrodynamic lengthscale |
| $\hat{D}_E$ | $D_E / (L^2 k_M^{\text{off}})$ | Diffusion of ECT-2 |
| $K_E^{\text{on}}$ | $k_E^{\text{on}} / (h k_M^{\text{off}})$ | Binding rate of ECT-2 |
| $K_{\text{ME}}$ | $k_{\text{ME}} M^{(\text{Tot})} / k_E^{\text{on}}$ | Relative recruitment ratio of ECT-2 by myosin |
| $K_{\text{AE}}$ | $k_{\text{AE}}$ | Inhibition of ECT-2 by AIR-1 |
| $K_E^{\text{off}}$ | $k_E^{\text{off}} / k_M^{\text{off}}$ | ECT-2 unbinding ratio |
| $\hat{A}_{\text{sat}}$ | $A_{\text{sat}} / A^{(\text{Tot})}$ | AIR-1 saturation level |
| $\hat{D}_M$ | $D_M / (L^2 k_M^{\text{off}})$ | Diffusion of myosin |
| $K_{\text{EM}}$ | $k_{\text{EM}} E^{(\text{Tot})} / (h k_M^{\text{off}})$ | Activation of myosin by ECT-2 (through Rho) |
| $K_{\text{fb}}$ | $k_{\text{fb}} (M^{(\text{Tot})})^k / k_M^{\text{off}}$ | Self-limiting myosin feedback |

### B.2 Numerical solution

We use standard numerical methods to solve (S1). We discretize the one-dimensional domain at *N* points with spacing 1*/*Δ*x*, and define the centered differentiation matrix ***D*** and standard three-point Laplacian differentiation matrix **Δ**. Given the myosin profile at time step *n*, we first compute the velocity *v*^(*n*)^ by solving *v*^(*n*)^ = *ℓ*^2^**Δ***v*^(*n*)^ + *ℓ****D****M* ^(*n*)^. Once the velocity is computed, the ECT-2 and myosin equations are solved by combining a first-order upwind finite volume scheme for the advection terms (Hundsdorfer et al., 2003, Sec. 1.4) with implicit treatment of the diffusion terms (using the standard three point Laplacian). The reaction terms are all treated explicitly, and time-stepping is first-order accurate.

### B.3 Parameter estimation

Parameter estimation for the dimensionless equations (S1) can be performed in three steps: first, we directly assign quantities that have already been measured experimentally. Second, we freeze the ECT-2 profile and assign parameters to the myosin equation (S4e) to match experimental data. Third, we choose the parameters in the ECT-2 equation (S4d) based on stability considerations.

#### B.3.1 Direct measurements

Some of the parameters can be determined directly from experimental measurements, as follows

1. The embryo cross section is an ellipse with approximate radii 27 *µ*m and 15 *µ*m, which gives a perimeter *L* = 134 *µ*m (Goehring et al., 2011).
2. The variable *ℓ* is the hydrodynamic length scale. In dimensional units, this was measured to be *ℓ* ≈ 13 *µ*m (Mayer et al., 2010), which means 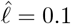 in (S1) (10% domain perimeter).
3. The myosin bound lifetime is about 15 s, according to measurements in the anterior of wild-type embryos, or in *par* mutant embryos which do not polarize (Gross et al., 2019, Fig. 1b). Since we typically model polarity *establishment*, where embryos are initially unpolarized, we set 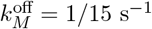.
4. We assume that all species have a dimensional diffusion coefficient *D*_*E/M*_ = 0.1 *µ*m^2^/s (Goehring et al., 2011; Gross et al., 2019; Robin et al., 2014). Rescaling length by *L* and time by 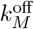 gives a dimensionless coefficient 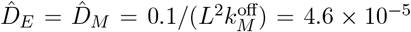. This dimensionless coefficient is sufficiently small as to render diffusion relatively unimportant in shaping the concentration fields. If we assume instead, for instance, that myosin cannot diffuse in the membrane, while ECT-2 has a ten-fold larger diffusion coefficient, the steady state profiles of ECT-2 and myosin are changed by at most 5% (see Fig. S6).
5. The ECT-2 lifetime was measured using FRAP to be on the order of a few seconds (Longhini and Glotzer, 2022, Fig. 3D). In cytokinesis, we set 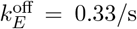, for a three second life-time. Rescaling gives 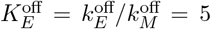. The data show slightly faster recovery during polarization, so we increase 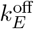 by 20% for those simulations.

**Figure S3:**
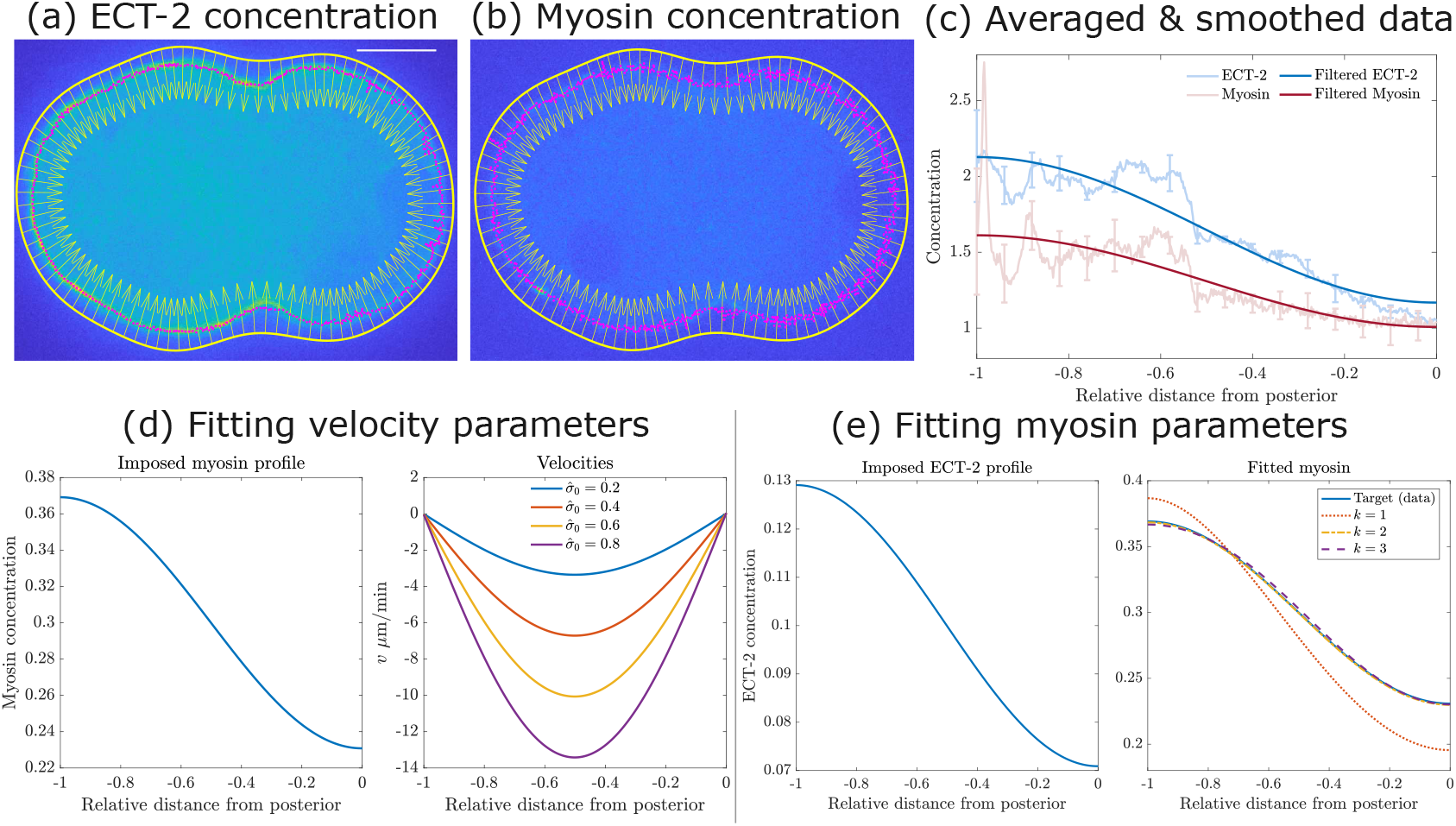
Fitting myosin and flow parameters from experimental data in wild-type embryos. (a–c) We extract an ECT-2 and myosin concentration from *N* = 3 ECT-2 mNG and myosin mKate embryos imaged during pseudo-cleavage. (a,b) Show embryo images, where the scale bar is 10 *µ*m, and (c) shows the resulting averaged and smoothed data over *N* = 3 embryos. (d–e) The ECT-2 and myosin profiles from the experimental data are used to obtain the parameters 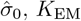, and *K*_fb_ for each *k*.

### B.2 Parameters for myosin equation

To infer the myosin parameters 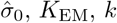 and *K*_fb_, our strategy is to impose the ECT-2 profile measured experimentally at the quasi-steady state of pseudo-cleavage, then solve for the myosin parameters required to match the experimental myosin profile. As shown in Fig. S3, we utilize previously-imaged embryos with ECT-2 and myosin reporters (Longhini and Glotzer, 2022, Fig. 1) for this purpose. For each set of images, we use Matlab’s built-in algorithms to segment the embryo boundary and compute an arclength parameterization of the boundary curve. Following this, we use Fourier filtering to filter the result and obtain a smoothed boundary with a normal vector at each point (see Fig. S3(a,b)). For each point on the arclength curve, we draw a 30 pixel (3 *µ*m) line inward and compute the maximum intensity along this line. Averaging over all frames where pseudo-cleavage is present, then repeating for three embryos to generate error bars, gives the curves shown in Fig. S3(c). Fitting these curves with a Fourier interpolant then gives smoothed representations that can be used for fitting. We note that the punctate myosin profile during establishment phase can somewhat confound the myosin measurements, but our data show a clear general trend that is captured by the Fourier interpolant.

To transition the target curves to model inputs, we scale by the expected amount of bound protein. Experimental data (Longhini and Glotzer, 2022, Fig. 1) show that about 10% of ECT-2 is bound to the cortex. This estimate is based on Fig. S3(c), which shows the average ECT-2 intensity in the cortical region to be about 1.5 times the cytoplasm. If the embryo is an ellipsoid with radii 27 × 15 × 15 *µ*m and the cortex has thickness 400 nm, then the cortex is 6.7% of the total volume. Multiplying by 1.5 gives 10% of the total ECT-2 bound. While the specific number (10%) is of little consequence to our model, the relative abundance of ECT-2 in the cytoplasm demonstrates that cytoplasmic depletion will not play a role in the dynamics. A similar set of data (Gross et al., 2019, Figs. S2,S3) show that about 30% of myosin is bound to the cortex. With these parameters, we scale the smoothed curves to obtain target curves for the model (the imposed ECT-2 curve is the experimentally-measured curve, but scaled to have mean 0.1).

To infer the velocity strength 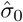 (Fig. S3(d)), we fix the myosin profile, then solve (S1c) to get the velocity profile and convert to *µ*m/min. Previous work (Gross et al., 2019, Fig. 3j) found the velocity in wild-type embryos to be at most 10 *µ*m/min. For this reason, we set 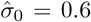 (Fig. S3(d)). Following this, we use the observed ECT-2 profile to set the remaining myosin parameters. We fix 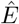 at the measured experimental profile, then solve the myosin equation (S4e) with three different values of *k* = 1, 2, 3 to fit *K*_EM_ and *K*_fb_ to match the target myosin profile. When *k* = 1, the solution to (S1) can be written as (neglecting diffusion and advection)

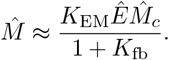

Since advection will only accentuate asymmetries, the minimum myosin asymmetry (max/min) that results from the imposed ECT-2 profile is 1.9 (equal to the ECT-2 asymmetry). Because the asymmetry in the myosin data (Fig. S3(c)) is 1.5, we need nonlinear inhibition (*k >* 1 in (S4e)) to match the myosin profile. As shown in Fig. S3(e), using *K*_EM_ = 9 and *K*_fb_ = 3.6 for *k* = 2, while *K*_EM_ = 6.5 and *K*_fb_ = 5.25 for *k* = 3 allows the solution of (S1) to match the smoothed experimental data.

### B.3 Fitting the ECT-2/AIR-1 parameters

There are now four parameters remaining: 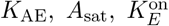, and *K*_ME_. The saturating AIR-1 threshold *A*_sat_ cannot be fit from polarization conditions because AIR-1 signals are low; as such we set its value according to the investigation in Appendix A (*A*_sat_ = 0.25). The parameter *K*_AE_ can be set by simulating AIR-1 activity in the absence of myosin. In myosin-depleted embryos, it was previously-shown that the ECT-2 asymmetry during early polarization is 1.2 (Longhini and Glotzer, 2022, Fig. 3A), which implies that the local AIR-1 activity induces a 20% depletion of AIR-1 on the posterior. We therefore infer *K*_AE_ by simulating polarization (as in Fig. 3) with no myosin activity (setting *K*_ME_ = 0 and 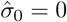). As shown in Fig. S4 (black lines), the steady state ECT-2 asymmetry is 1.2 when *K*_AE_ = 1.3.

With the AIR-1 parameters set, there is effectively one parameter remaining: the strength of indirect recruitment of myosin on ECT-2 (there is also 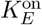, the basal ECT-2 binding rate, but this is set to maintain 10% bound ECT-2 on the cortex). We take a systematic approach to setting this parameter: as shown in Fig. S4, setting *K*_ME_ = 0, so that the only interaction of ECT-2 with myosin comes via flows, gives a negligible change in the ECT-2 profile from the AIR-1-only case (no myosin). Thus, indirect recruitment must be responsible for shaping the ECT-2 profile. To fit a value, we increase *K*_ME_ until we trigger locally oscillatory patterns of ECT-2 accumulation. As shown in Fig. S4, these patterns occur upon the transition to the regime where the uniform ECT-2 profile is linearly unstable (see the next section for an analysis). The values we use are *K*_ME_ = 2.5 (for *k* = 2) or *K*_ME_ = 1.5 (for *k* = 3).

#### B.3.4 The parameters are on the edge of the stability boundary

To perform linear stability analysis of the model equations (S1) with *K*_AE_ = 0, we perturb the myosin and ECT-2 profiles around the uniform state by setting *M* = *M*_0_ + *δM* and *E* = *E*_0_ + *δE*, where *δM* = Σ_*j*_ *m*_*j*_(*t*) exp (2*πijx*), and likewise for *δE*. Substituting this representation for *M* into the velocity equation (S1c) then gives a representation for the velocity *v* =Σ_*j*_ *v*_*j*_(*t*) exp (2*πijx*), where (Bois et al., 2011, Eq. (11))

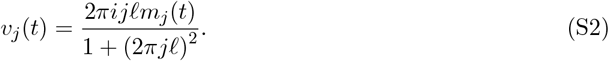

**Figure S4:**
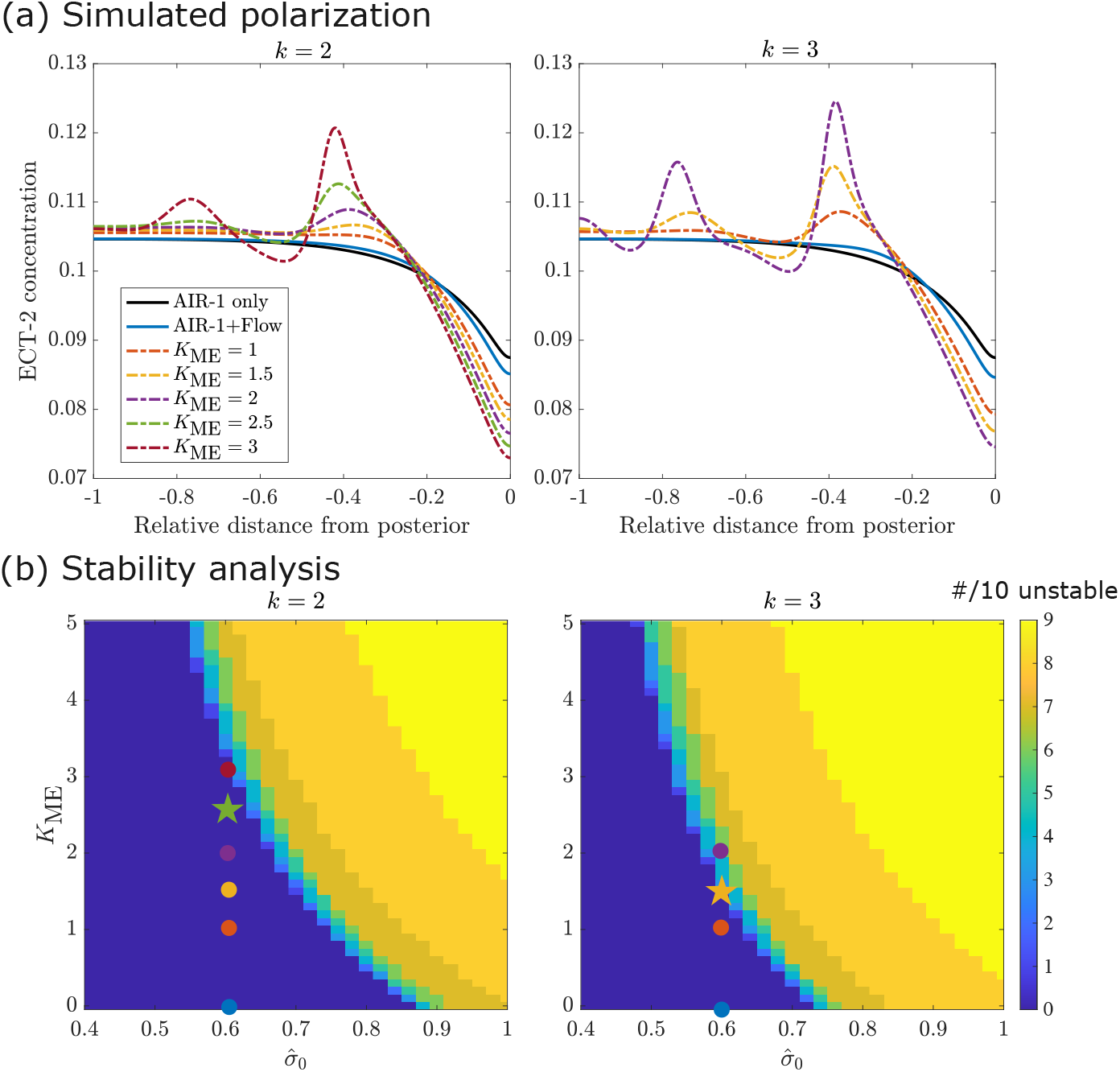
Fitting the ECT-2 parameters, and relating those to the stability boundary. Top plots: for each *k*, we simulate polarization using the AIR-1 cue from centrosomes 1.9 *µ*m away from the cortex. The black lines show the ECT-2 profile without myosin, while the blue lines show the ECT-2 profile with advection only (no indirect recruitment). Other colored lines show the ECT-2 profile with advection *and* indirect recruitment by myosin. Bottom plots: the stability diagram as a function of 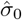 (flow strength) and *K*_ME_ (indirect recruitment strength). The chosen parameters (stars) lie on the edge of the unstable regime. In all cases, 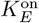 is set so that 10% of ECT-2 is bound to the cortex.

Substituting this representation into the ECT-2 and myosin equations, and keeping terms to linear order in *δ* gives the matrix equation

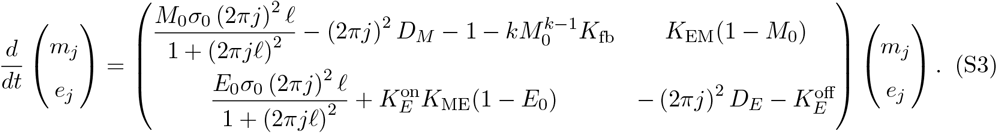

The dynamics are unstable if the 2×2 matrix above has a positive eigenvalue (negative determinant). The stability diagram in Fig. S4 shows the number of unstable modes for each pair 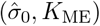, with all other parameters fixed to their default values (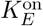 is adjusted to maintain 10% ECT-2 bound). The most unstable behavior (quantified by how many of the first 10 Fourier modes *j* = 1, … 10 in (S3) are unstable) occurs for high flow speeds and high recruitment rates.

### B.4 Computational experiments

This section presents results on some computational experiments motivated by the main text.

#### B.4.1 Nonlinear negative feedback

We first show (Fig. S5) that the steady state simulated ECT-2 profiles during polarization and cytokinesis are unchanged when we switch the nonlinear negative feedback exponent *k* in (1b) from *k* = 2 (main text) to *k* = 3 (Fig. S5). Comparing the polarization (Fig. 3(d)) and cytokinesis (Fig. 5) simulations, we find that the match between model and experiment is similar. The only small difference is that the *k* = 3 simulations have slightly stronger nonlinear negative feedback, and produce smoother profiles in cytokinesis when the AIR-1 signal is stronger (compare dark lines in the cytokinesis curves in Figs. 5 and S5).

#### B.4.2 Changing diffusion coefficients

During polarity establishment, myosin forms large foci which look stationary in the cell cortex, whereas ECT-2 is a more mobile protein. To examine if changing diffusion coefficients affects the steady state profile, in Fig. S6 we compare control diffusion coefficients (*D*_*E*_ = *D*_*M*_ = 0.1 *µ*m^2^/s to an alternative set where myosin is immobile (*D*_*M*_ = 0 *µ*m^2^/s) and ECT-2 is highly mobile (*D*_*E*_ = 1 *µ*m^2^/s). As shown in Fig. S6, the resulting steady state profiles of ECT-2 and myosin are similar in the two cases.

**Figure S5:**
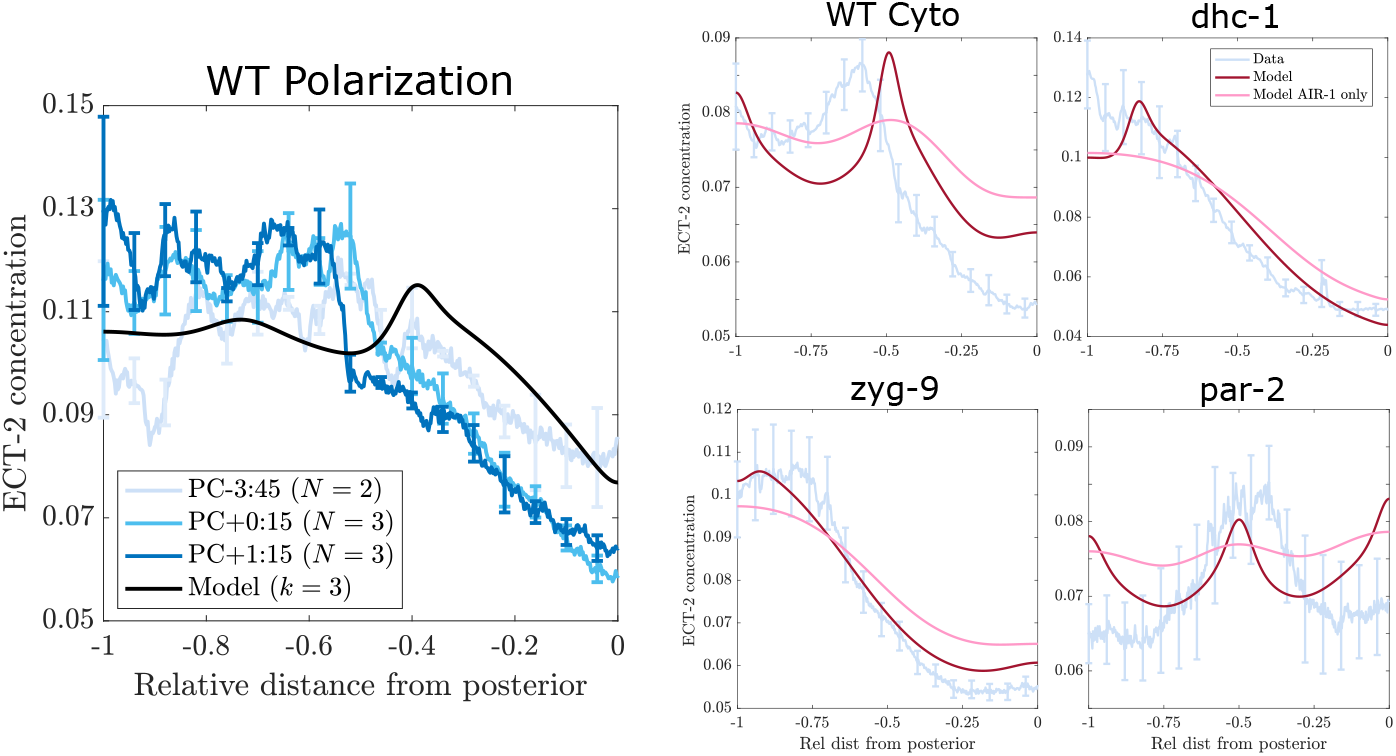
Representative results when *k* = 3 in the myosin equation (1b). We show the resulting ECT-2 profiles in WT polarization (c.f. Fig. 3(d)) and cytokinesis (c.f. Fig. 5)

**Figure S6:**
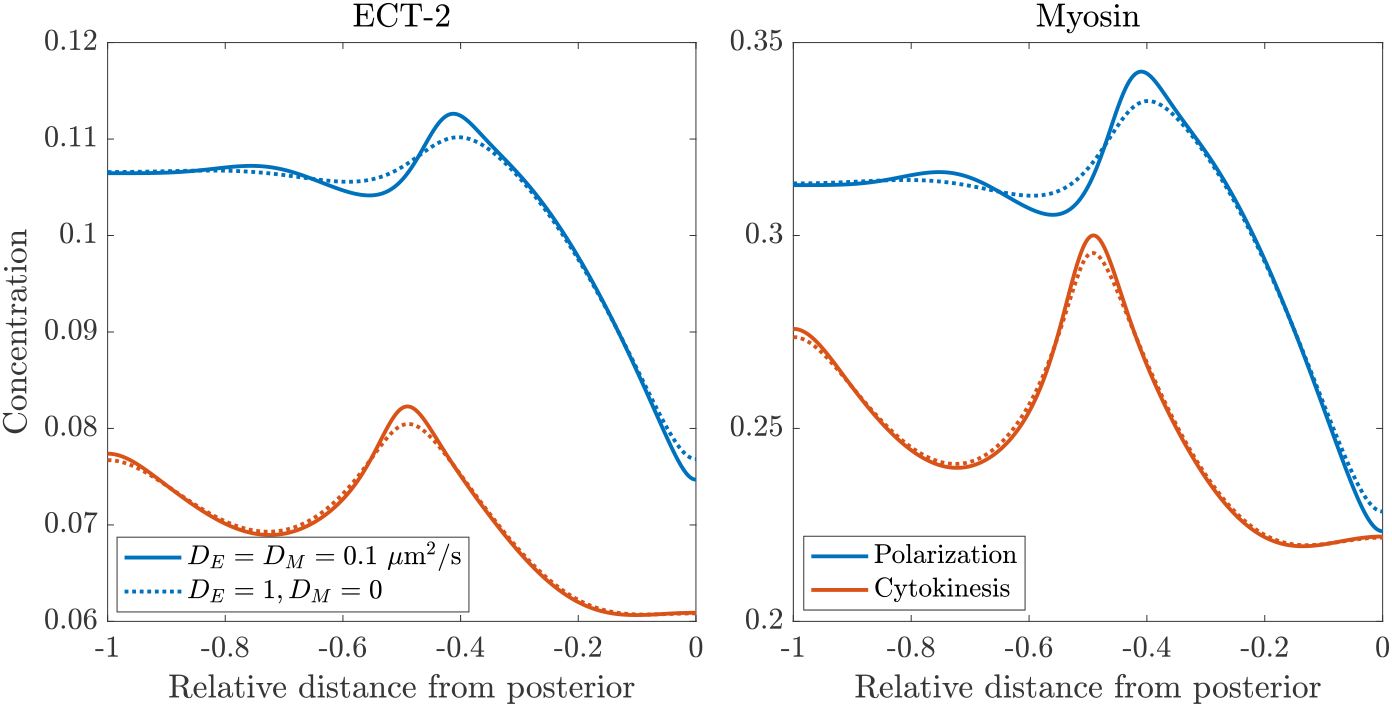
Steady states for polarization and cytokinesis in wild-type embryos assuming (solid lines) *D*_*E*_ = *D*_*M*_ = 0.1 *µ*m^2^/s (control conditions), and (dotted lines) *D*_*E*_ = 1 *µ*m^2^/s and *D*_*M*_ = 0. Blue lines show polarization, red lines show cytokinesis.

### B.4.3 Incorporating an explicit intermediary

To more thoroughly examine our hypothesis that “myosin” recruits ECT-2, we consider an explicit model where a third species (“P”) is advected with myosin and recruits ECT-2. The dimensional equations governing this situation are

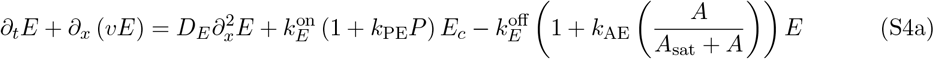

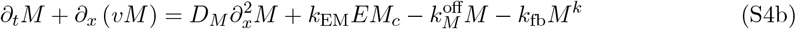

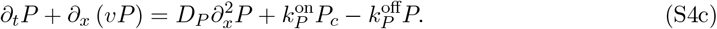

Similar to (S1), we nondimensionalize the equations to obtain the system

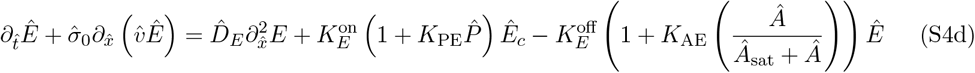

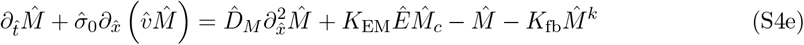

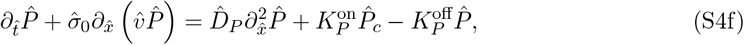

where 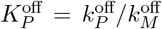, the binding rate 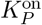 is arbitrary (we set it such that 30% of protein is bound, similar to myosin), and all parameters are the same as previously. In Fig. S7, we plot the corresponding steady ECT-2 profiles that result during polarization under different values of 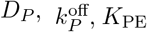. To advect *P* with cortical flows, we consider residence times similar to that of the longest-lived PAR proteins, 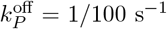 and 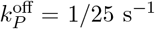 (Robin et al., 2014; Illukkumbura et al., 2023). Similar to the more minimal model (Fig. S4), a small amount of recruitment propagates AIR-1- and flow-driven asymmetries, but larger coefficients lead to instabilities and oscillatory profiles. More specifically, we find that this model of recruitment through an advected intermediary typically undergoes instabilities at smaller levels of posterior clearance than the minimal model (1) which utilizes recruitment by species that colocalize with myosin. Nevertheless, the results for high diffusivities and intermediate residence times of the intermediary (bottom right plot in Fig. S7) are quite similar to the model we consider in the main text.

#### B.4.4 Reproducing longer timescales of posterior myosin depletion

As discussed in the main text, the transient cue simulation in Fig. 4 is unphysiological because it involves a sudden removal of the centrosome cue at *t* = 5 mins. In Fig. S8 we attempt to correct for this by relaxing some of the model assumptions. First, we consider a simulation where the myosin exchange rates are four-fold slower (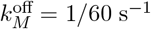 with *K*_EM_ fixed). To position the simulation at a similar point in the stability plane, we also modify the nonlinear negative feedback *K*_fb_ = 12, with all other (dimensional) parameters fixed. In this simulation (red curve in Fig. S8), the posterior depletion of ECT-2 and myosin recovers more slowly after the cue is removed at *t* = 300 s (30 s to half max instead of 15 s), but the recovery (driven by immediate recovery of ECT-2 at the posterior) is still quite fast. As a second possibility, we consider moving centrosomes, which sit 2 *µ*m from the cortex over the first five minutes, then shift linearly to 10 *µ*m at *t* = 10 mins, with the corresponding change in the AIR-1 signal (see Fig. 3(b)). While ECT-2 still responds rapidly to changes in the AIR-1 signal, the slow changes in the AIR-1 signal drive linear posterior recovery of ECT-2 (yellow curve), which accords better with the experimental data.

**Figure S7:**
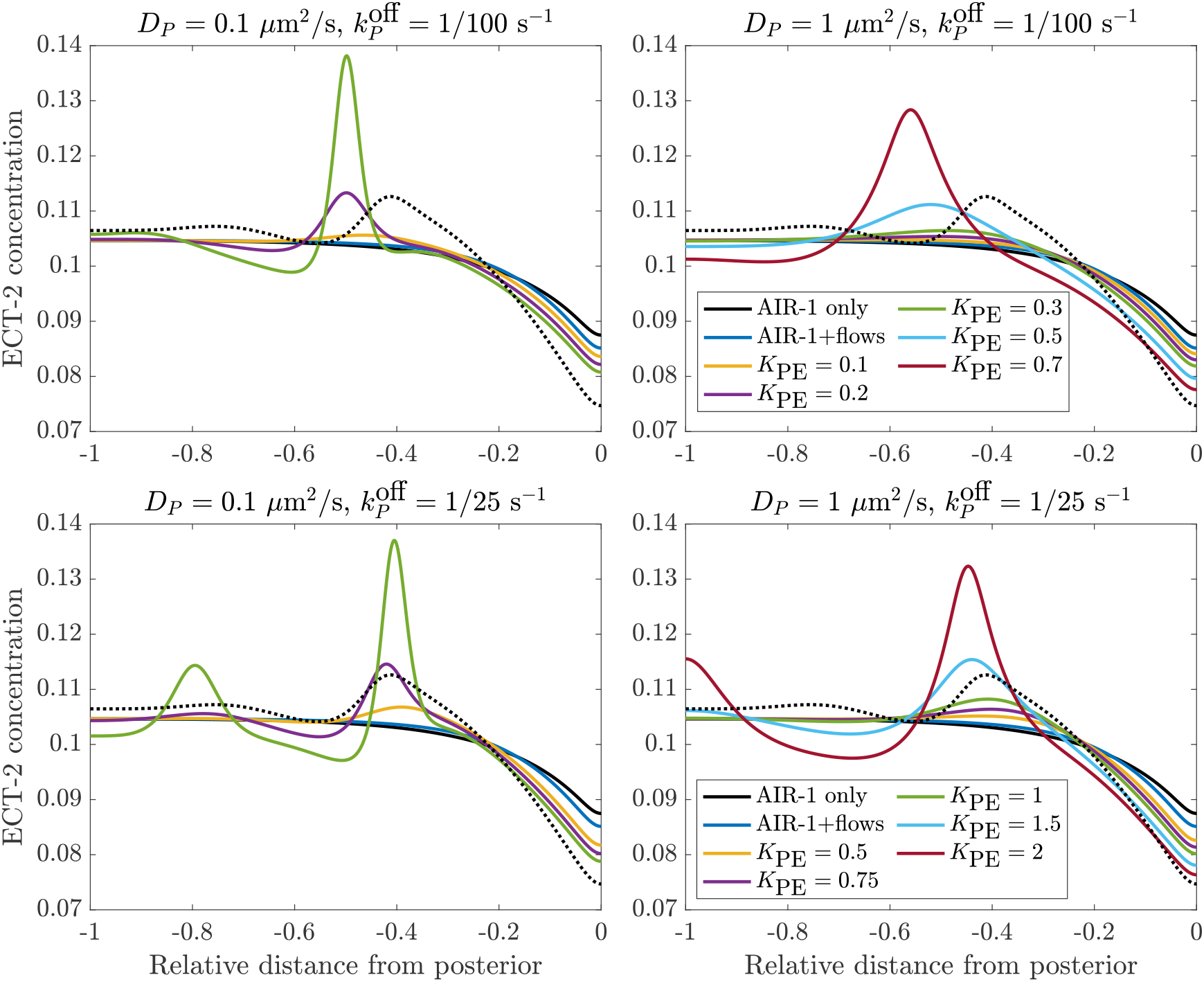
Simulating polarization using the modified model (S4) where a longer-lived protein recruits ECT-2. Solid lines show the results from the recruitment model (S4) with the indicated parameters, while the dotted black line is the steady state for the “minimal model” model (1) in the main text.

**Figure S8:**
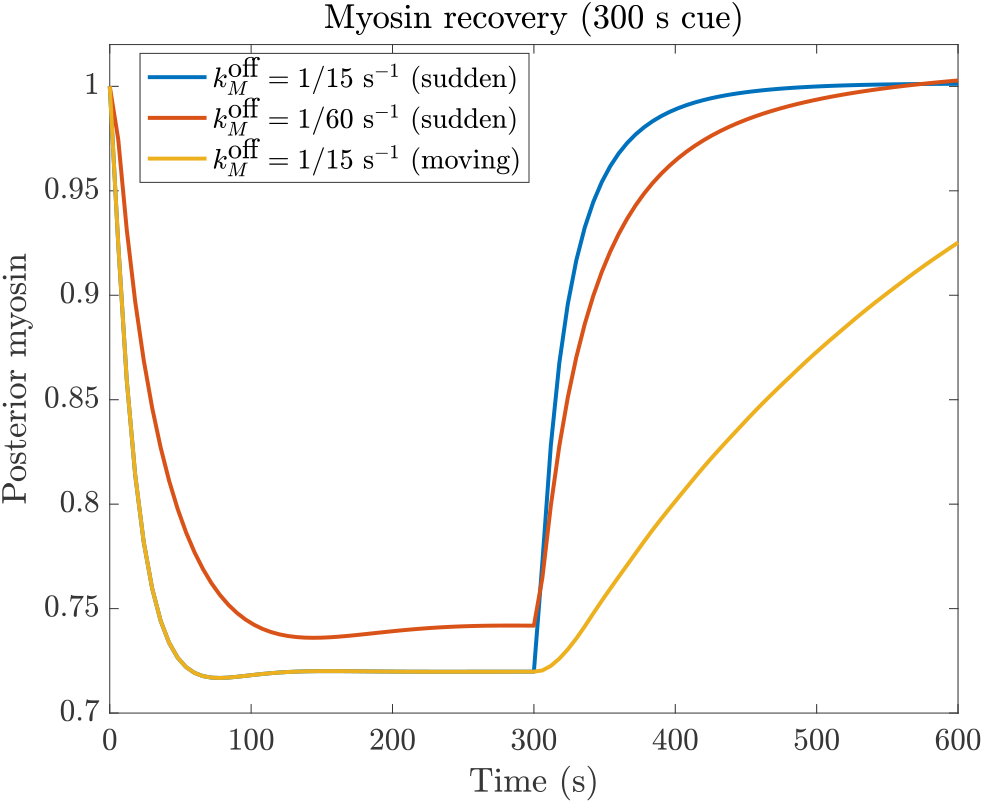
Simulating myosin recovery during transient polarization. We show the myosin concentration at the posterior pole (normalized by the steady state) under three different conditions: control (blue, where the AIR-1 cue is present for 5 minutes), a simulation where the myosin lifetime is four times longer (red), and a simulation where, starting at *t* = 5 mins, the centrosomes move at a linear rate from 2 to 10 *µ*m from the cortex (yellow, instead of the cue disappearing suddenly).

#### S.4.5 AIR-1 and myosin depletion during cytokinesis

To further probe the model during cytokinesis, we compare the output upon depletion of AIR-1 or myosin. In Fig. S9, we begin with AIR-1 depletion at left, showing the measured ECT-2 accumulation at cleavage onset in wild type (solid line) and AIR-1(RNAi) (dotted line). Using the same normalization between the two curves, we observe more ECT-2 accumulation globally under AIR-1 depletion, and a gradient that is less steep. The profile under AIR-1 depletion can be reproduced in the model by multiplying the wild-type AIR-1 signal (shown in Fig. S2) by 0.25 (to mimic 75% depletion). Further depletion causes gradients in ECT-2 accumulation to vanish.

**Figure S9:**
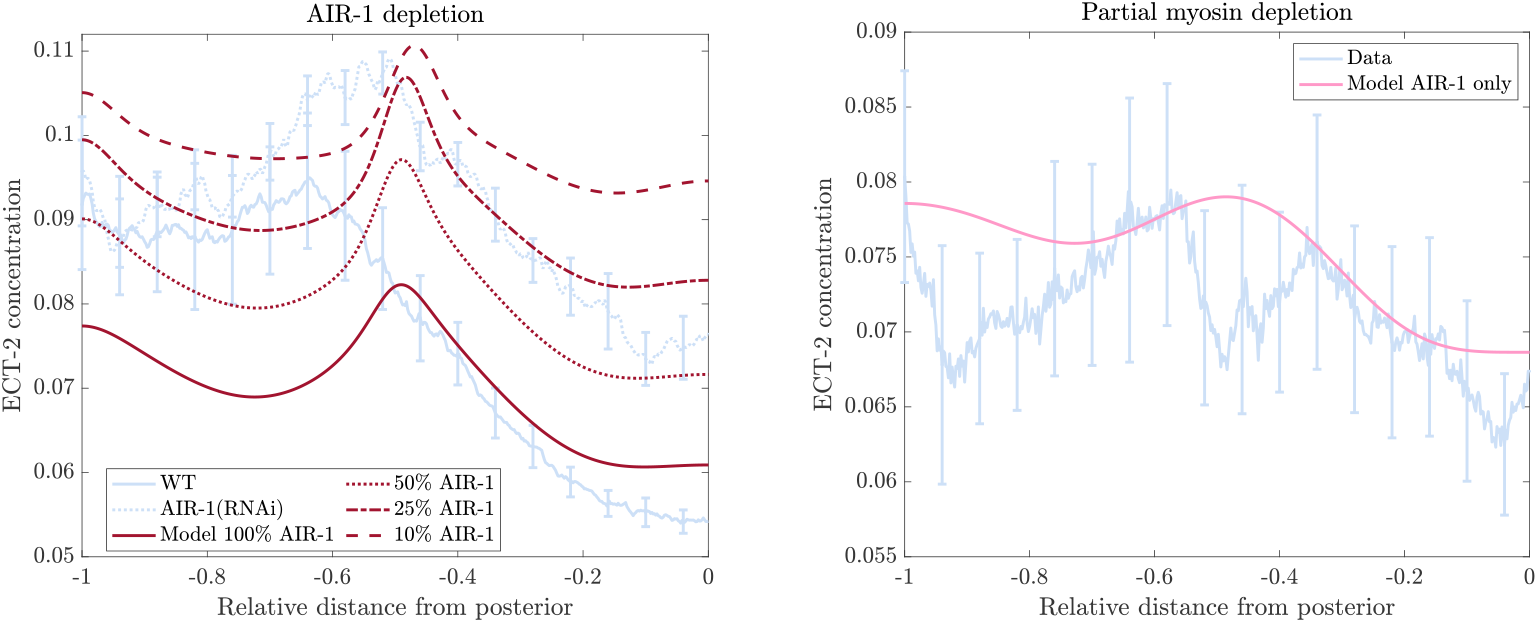
Steady state ECT-2 accumulation during cytokinesis under AIR-1 and myosin depletion. Left plot: we show the steady state accumulation in the 50 s following cleavage ingression in control (wild type; *N* = 9) and AIR-1(RNAi) embryos (*N* = 11) (the constant of normalization is the same in both cases). We compare this to the model’s simulated steady state using the wild-type AIR-1 profile shown in Fig. S2, multiplied by 1 (solid line), 0.5 (dotted line), 0.25 (dashed-dotted line), and 0.1 (dashed line). Right plot: we compare the AIR-1 only model prediction (pink line, same as the left panel of Fig. 5) to experimental data where myosin is partially depleted (*N* = 8; blue lines), again looking at the 50 s time interval that follows cleavage ingression.

To further validate that the relative effects of AIR-1 and myosin are similar in polarization and cytokinesis, in Fig. S9 we also consider experimental data for embryos under partial myosin depletion. We plot the experimental profile using a normalization constant taken from control embryos, then compare the result to the model with no myosin-based flows or indirect recruitment (pink line repeated from the left panel of Fig. 5; this uses the centrosome positions in wild-type embryos). The experimental data and model both predict near-uniform ECT-2 accumulation, suggesting that the effect of AIR-1 alone on ECT-2 (parameter *K*_AE_ previously set to match polarization) is correct during cytokinesis as well.

## Notes

https://github.com/omaxian/CElegansModel/

### Competing Interest Statement

The authors have declared no competing interest.

### Summary of Updates

The manuscript was revised in response to reviewers comments. The modeling evolved somewhat and the manuscript was extensively edited.

### Funding

- National Institute of General Medical Sciences: 5R35GM127091

